# Hypoxia-induced metastatic heterogeneity in pancreatic cancer

**DOI:** 10.1101/2025.08.26.672389

**Authors:** Pradeep Moon Gunasekaran, Qianqian Wang, Yoke-Chen Chang, Polina Guseva, Rajika Chauhan, Alexander Kley, Gene Lee, Siddharth Ghosh Roy, Yousef Masoudpoor, Arthur Roberts, Kelly Watkins Walton, Lucyann Franciosa, Shafiq Bhat, Emmanuel Zachariah, Kishan Patel, Zhongren Zhou, Wenjin Chen, Julie Zhouli Ni, Sam Guoping Gu, Cristina Montagna, Shin-Heng Chiou

**Affiliations:** Rutgers Cancer Institute, New Brunswick, New Jersey, USA; Department of Medicine, Rutgers Robert Wood Johnson Medical School, Rutgers the State University of New Jersey, New Brunswick, New Jersey, USA; The College of New Jersey, Ewing Township, New Jersey, USA; Department of Molecular Biology and Biochemistry, Rutgers the State University of New Jersey, Piscataway, New Jersey, USA; Department of Biological Sciences, Rutgers the State University of New Jersey, New Brunswick, New Jersey, USA; Department of Pathology and Laboratory Medicine, Rutgers Robert Wood Johnson Medical School, Rutgers the State University of New Jersey, New Brunswick, New Jersey, USA

## Abstract

In most solid tumors, hypoxia constitutes a defining microenvironmental feature that reprograms malignant cells into a highly metastatic state by driving cellular plasticity and exacerbating chromosomal instability (CIN). However, the mechanisms by which cancer cells concurrently co-opt these elements of hypoxic adaptation to promote metastasis remains poorly understood. Here, we report that hypoxia promotes metastasis by suppressing the JmjC-containing histone lysine demethylase Kdm8. CRISPR/Cas9-mediated targeting of *Kdm8* in a *Kras*;*Trp53*-driven mouse model of pancreatic ductal adenocarcinoma (PDA) robustly rewires the malignant cell transcriptomic programs, leading to a profound loss of the epithelial morphology and widespread metastatic disease. In PDA patients, a high KDM8-induced gene signature is associated with reduced metastatic burden and better survival in advanced disease. Notably, *Kdm8* suppression in normoxia recapitulates key aspects of the global epigenetic and transcriptomic rewiring, mitotic spindle defects, and CIN induced by hypoxia. Moreover, disruption of Kdm8’s demethylase activity phenocopies *Kdm8* loss, whereas expression of hypermorphic Kdm8 variants resistant to hypoxic suppression markedly reduces metastasis beyond the levels achieved by the wildtype protein. Through the suppression of Kdm8 demethylase function, hypoxia unleashes a potent metastatic program by simultaneously advancing cellular plasticity and CIN.

## Introduction

Pancreatic ductal adenocarcinoma (PDA) remains one of the deadliest malignancies, largely due to its highly aggressive metastatic behavior ^1^. As with other carcinomas, PDA metastasis requires the acquisition of diverse and often opposing cellular programs that enable dissemination and colonization at distant sites ^2^. In line with this, paired assessments of genomic alterations in primary and metastatic tumors provided limited insights into the molecular drivers of metastasis ^3–5^. In contrast, recent studies highlight transcriptomic and epigenetic plasticity as key malignant features underlying treatment resistance, subtype differentiation, and metastatic progression in PDA ^6–17^. Despite these advances, the cancer cell-extrinsic factors within the tumor microenvironment that drive metastatic plasticity remain elusive.

Intratumoral hypoxia is a potent inducer of metastasis ^18^. Under hypoxic conditions, the stabilization of the hypoxia-inducible factors (HIFs) promotes epithelial-to-mesenchymal transition (EMT), a gene expression program broadly linked to metastasis across multiple cancer types ^19^. In PDA specifically, EMT has been shown to facilitate the metastatic dissemination ^6,12,20,21^. However, genetic depletion of *Hif1a*, which encodes the hypoxia-stabilized subunit of the Hif1a/Arnt heterodimer, unexpectedly accelerated Kras^G12D^-driven pancreatic tumor progression and metastasis ^22,23^, suggesting that hypoxia may act through additional, Hif1-independent mechanisms. Beyond HIFs signaling, hypoxia triggers extensive epigenetic reprogramming, in part by inhibiting the activity of JmjC-domain-containing histone lysine demethylases (KDMs), which require oxygen (O₂) as a cofactor ^24,25^. In human PDA, genetic loss of *KDM6A* (∼6%), an X-linked histone 3 lysine 27 (H3K27)-specific demethylase, is associated with the basal-like or squamous-like state and poor prognosis ^26–28^. Yet, studies using genetically engineered mouse (GEM) models indicate that Kdm6a suppresses PDA progression through a demethylase-independent mechanism, raising the possibility that other KDM members may mediate hypoxia-driven reprogramming of the malignant state ^26^. In addition to its epigenetic effects, hypoxia is known to promote metastasis through DNA overreplication ^29^. Hypoxia-induced DNA damage and genomic alterations have traditionally been attributed to the suppression of multiple DNA damage response pathways ^30^. Large genomic studies utilizing bulk whole genome sequencing (WGS) data from patients suggest a clear link between hypoxia in solid tumors and distinct mutational and copy number alteration (CNA) signatures ^31,32^. Despite this, the mechanism by which hypoxia promotes chromosomal instability (CIN) remains unclear ^30^. CIN in cancer enables the selective amplification of genomic loci that enhance cancer cell fitness under stress conditions, including hypoxia ^33–37^. One such examples is *KRAS* copy number gain, which becomes increasingly prevalent during PDA progression and is most pronounced in metastatic lesions when compared to the primary site ^38–40^. Mounting evidence suggests that oncogenic *KRAS* drives PDA progression through diverse mechanisms, including metabolic re-adaptations ^41–44^, immune suppression ^45–47^, and subtype differentiation ^39,48^. However, whether and how *KRAS* copy number in PDA arises as a direct consequence of hypoxia remains unresolved.

KDM8 is a histone lysine demethylase that targets H3K36 di-methylation (H3K36me2) and regulates cell proliferation in mammalian cells ^49,50^. In *C. elegans*, disruption of the *KDM8* ortholog *jmjd-5* results in the loss of germ cell identify and reduced fertility ^51^. Genetic depletion of *Kdm8* in mice leads to embryonic lethality, attributed to dysregulation of *Cdkn1a* via increased H3K36me2 levels ^50,52^. In human embryonic stem cells, *KDM8* (a.k.a. *JMJD5*) is required to maintain pluripotency ^53^, and germline mutations are associated with severe developmental disorders including intellectual disability and craniofacial abnormalities ^54^. Furthermore, jmjd-5/KDM8 has also been implicated in regulating chromosome segregation and maintaining genomic integrity, presumably through its JmjC domain-dependent activity ^55,56^. Despite its established roles in development, pluripotency, and genome maintenance, the role of KDM8 in regulating cellular plasticity and metastatic progression in PDA remains unknown.

In this study, we present compelling evidence that hypoxia promotes PDA metastasis through the suppression of KDM8. Using a novel GEM model that employs CRISPR/Cas9 genome editing and barcoded *Kras^G12D^* alleles ^57,58^, we show that *Kdm8* depletion reprograms the epithelial state toward EMT, driving widespread metastasis. Notably, the metastatic potential of PDA can be modulated by toggling the activity of KDM8’s JmjC domain. These findings offer new mechanistic insight into how hypoxia enhances the metastatic proclivity of malignant PDA cells.

## Results

### Knockdown of Kdm8 drives metastasis in multiple transplanted tumor models

Unlike the HIF pathway, select KDM family members respond to hypoxia by inducing immediate epigenetic changes ^24^. Our initial effort focused on identifying functionally important KDM family members that (i) induce embryonic lethality when genetically deleted and (ii) play a crucial role in regulating PDA metastasis. We performed subcutaneous tumor studies using the immunocompromised *NSG* (*NOD-scid;Il2rg^null^*) mice to functionally interrogate five prioritized *Kdm* genes for their effect on lung metastasis using the 688M *Kras^G12D^*;*Trp53^R172H^* murine PDA cell line (Figures 1A, S1A, and S1B) ^11^. Compared to the other *Kdm* family members, *Kdm8* knockdown with two independent short hairpin RNAs (shRNAs) reproducibly promoted lung metastasis without affecting subcutaneous primary tumor growth (Figures 1B and S1C-S1E).

**Fig. 1.**
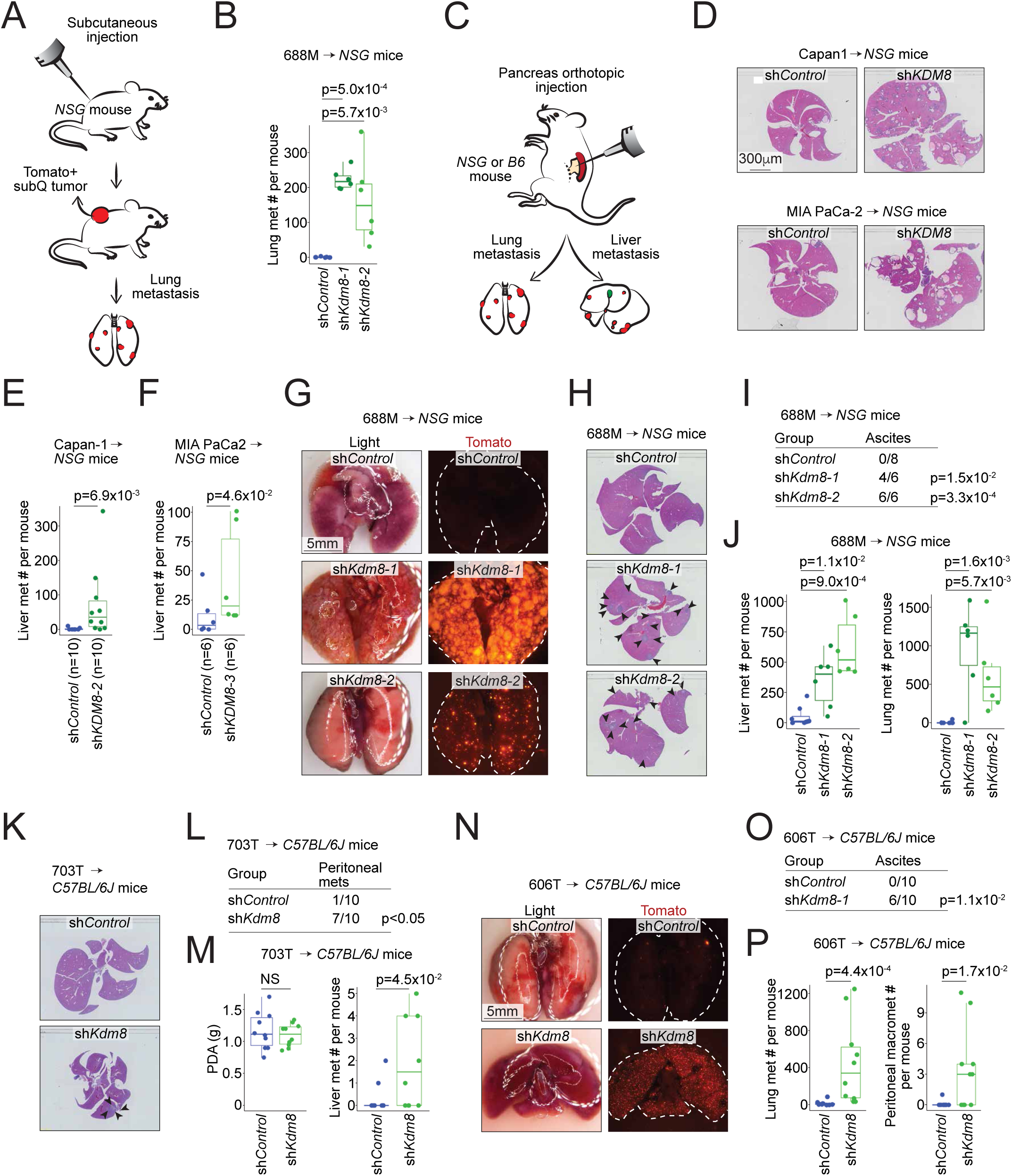
*KDM8* is a PDA metastasis suppressor gene. (**A**) Schematic of the subcutaneous tumor model for the evaluation of lung metastasis using the Tomato-positive 688M PDA cell line. (**B**) Lung metastasis counts in the *NSG* mice subcutaneously transplanted with 688M cells expressing control (sh*Control*, n=8) or shRNAs targeting *Kdm8* (sh*Kdm8-1*, n=6; sh*Kdm8-2*, n=6). The experiments were repeated at least three times with similar results. (**C**) Schematic of the pancreatic orthotopic tumor model for the evaluation of lung and liver metastasis. (**D**) Representative haematoxylin and eosin staining (H&E) of the livers from *NSG* mice receiving sh*Control* or sh*KDM8*-expressing Capan-1 (top) and MIA PaCa-2 (bottom) human PDA cells. (**E**,**F**) Liver metastasis counts per *NSG* mouse orthotopically transplanted with Capan-1 (**E**) or MIA PaCa-2 (**F**) cells expressing sh*Control* or sh*KDM8*. p-values were derived using Wilcoxon test. The experiments were repeated twice with similar results. (**G**-**J**) Representative light (left) and fluorescent (right) images of the lungs (**G**) and H&E staining of the livers (**H**), tallies of mice with ascites (**I**), and counts of the liver (left, **J**) and lung (right, **J**) metastases in the *NSG* mice orthotopically transplanted with 688M PDA cells expressing sh*Control* or two sh*Kdm8*s as in (B). The experiments were repeated three times with similar results. (**K**-**M**) Representative H&E of the livers (**K**), tallies of mice with peritoneal metastases (**L**), primary PDA tumor weight (left, **M**), and counts of liver metastases in the *C57BL/6J* (*B6*) mice (right, **M**) orthotopically transplanted with the Tomato-negative 703T PDA cells expressing sh*Control* or sh*Kdm8*. (**N**-**P**) Representative light (left) and fluorescent (right) images of the lungs (**N**), tallies of mice with ascites (**O**), and counts of the lung (left, **P**) and peritoneal (right, **P**) metastases in the *B6* mice orthotopically transplanted with the Tomato-positive 606T PDA cells expressing sh*Control* or sh*Kdm8*. The experiments were repeated twice with similar results. NS, not significant. Each dot is a mouse and boxes represent medians with interquartile range between 25th and 75th percentiles. Arrowheads indicate liver metastases.

To further investigate the role of KDM8, we orthotopically transplanted control and sh*KDM8*-expressing human PDA cells, Capan-1 and MIA PaCa2, which express high levels of endogenous KDM8 protein, into the *NSG* mice (Figures 1C and S1F). Consistent with the subcutaneous tumor experiments, *KDM8* knockdown promoted liver metastasis in these orthotopic PDA tumor models (Figures S1G-S1J, and 1D-1F). Furthermore, compared to the control counterpart, orthotopically transplanted *Kdm8*-deficient 688M tumors seeded between 12 and 133-fold more metastases in the livers and lungs, respectively (Figures 1G-1J and S1K). Notably, the increased metastases were accompanied by more frequent ascites formation in mice transplanted with *Kdm8*-deficient 688M cells (Figure 1I). Similarly, *Kdm8* knockdown in two other murine PDA cell lines (703T and 606T, both *Kras^G12D^*;*Trp53^flox/flox^*) greatly promoted ascites formation and metastatic burdens in the lungs and livers when transplanted orthotopically into the immunocompetent *C57BL/6J* mice (Figures 1K-1P and S1L-S1O). Comparable to the findings in *NSG* mice, *Kdm8*-deficient 606T orthotopic tumors seeded 25-fold more pulmonary micrometastases compared to the control group, with many metastatic lesions appearing as individual cancer cells (Figures 1P and S1N). In summary, we screened five KDM family members and identified *Kdm8* as a potential metastasis suppressor in PDA. Rigorous transplanted tumor studies employing multiple *in vivo* models demonstrated that *Kdm8* and *KDM8* knockdown in both murine and human PDA cell lines, respectively, increased PDA metastasis. These results were consistent across various transplantation routes, the *Trp53* status of the murine PDA cell lines, short hairpin sequences for *Kdm8* targeting, and the presence or absence of anti-tumor immune surveillance.

### Establishing a novel autochthonous PDA mouse model that enables Kdm8 genetic depletion using somatic genome editing

To functionally interrogate *Kdm8* in neoplasms initiated by oncogenic *Kras* and *Trp53* inactivation, we built upon a previous approach ^58^ that enables the induction of spontaneous PDA in the *Trp53^flox/flox^*;*R26^LSL-Tomato^*;*H11^LSL-Cas^*^9^ mice (*PTC*; Figure 2A) by delivering an engineered adeno-associated virus (AAV) into the adult murine pancreas through a retrograde pancreatic ductal injection procedure ^57^. The engineered AAV for PDA initiation includes a U6 promoter-driven expression of an sgRNA targeting exon2 of *Kras* (U6-sg*Kras*), a 2.1Kb *Kras* genomic template with the codon 12 glycine-to-aspartic acid mutation (*Kras^G12D^*) flanked by diverse combinations of synonymous mutations that serve as molecular barcodes for lineage tracing and a set of sg*Kras*-resistant synonymous mutations, and PGK-*Cre* (Figure 2B). Pancreatic infusion of the engineered AAV into adult mice leads to the expression of sg*Kras* and Cre, which in turn drives the expression of Cas9, deletion of *Trp53*, and introduction of the engineered *Kras* variant into the *Kras* locus through homology-directed repair, ultimately facilitating the growth of PDA that is histologically indistinguishable from those derived from conventional *KPC* (*Kras^LSL-G12D^*;*Trp53^LSL-R172H^*or *Trp53^flox/flox^*;*Pdx1-Cre*) mice ^57^. Following the surgical procedure, an overall 74.2% of the mice developed histologically confirmed PDA by the time endpoints were reached (n=23/31, Table S1). Two cohorts received AAV engineered to carry either an additional U6-sg*Kdm8* (*PTC^Kdm8KO^*, n=18) or non-targeting U6-sg*NT* (*PTC^NT^*, n=13) cassette. To assess the metastatic burdens in these mice, 23 *PTC* mice with locally advanced PDA were carried forward to subsequent analyses (*PTC^NT^*, n=13; *PTC^Kdm8KO^*, n=10). Similar to the control cohort, *PTC^Kdm8KO^* mice developed solid tumors 4-6 months post-surgery with comparable overall survival (Figures 2C and 2D). For each *PTC^Kdm8KO^* tumor (total n=7), we detected a minimum of 80% indel (insertion and deletion) formation at the *Kdm8* targeted site estimated by the Surveyor nuclease assay, thus validating the effectiveness of the current method in deleting *Kdm8* (Figure 2E).

**Fig. 2.**
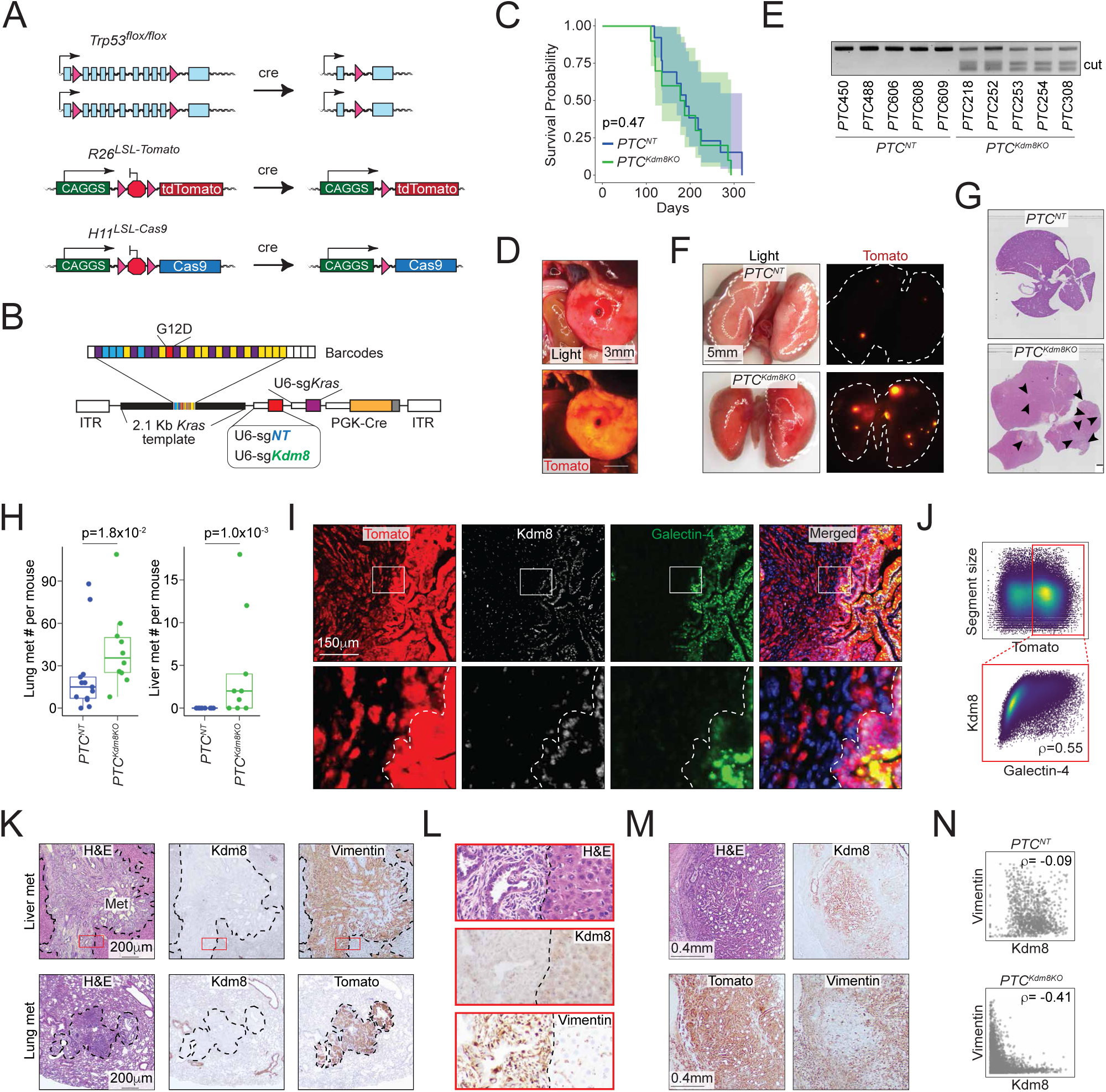
CRISPR-mediated *Kdm8* mutation induces profound loss of differentiation and widespread metastatic lesions in autochthonous PDA. **(A)** Schematic of the allelic configurations prior to and post Cre-induced recombination in the *PTC* mice. (**B**) Schematic of the pooled AAV library. Barcodes represent codons 2-31 of the murine *Kras*, engineered with two sets of synonymous (wobble) mutations (purple and yellow), along with the non-synonymous *Kras^G12D^* mutation (red) and four fixed synonymous mutations (blue) that confer resistance to sg*Kras* (purple). The engineered coding sequence is flanked by ∼2.1Kb *Kras* homology arms (black). sg*Kras* encodes an sg*RNA* targeting *Kras*. An additional *U6*-driven sg*RNA* - either a non-targeting (*U6*-sg*NT*) or *U6*-sg*Kdm8* - is included to serve as a control or to target *Kdm8*, respectively. *ITR*, inverted terminal repeats. (**C**) Kaplan-Meier survival analysis of *PTC* mice receiving AAV-sgNT (*PTC^NT^*) or AAV-sg*Kdm8* (*PTC^Kdm8KO^*). p = 0.16 using log rank test. (**D**) Representative light and fluorescent images of a *PTC^Kdm8KO^* tumor. (**E**) Agarose gel electrophoresis of genomic DNA from control and *PTC^Kdm8KO^* malignant cells using Surveyor nuclease assay. (**F**) Representative light and fluorescent images of the lungs from a control and *PTC^Kdm8KO^* mouse. (**G**) Representative haematoxylin and eosin staining (H&E) of the livers from a *PTC^NT^*and *PTC^Kdm8KO^* mouse. (**H**) Counts of lung (left, *PTC^NT^*, n=13; *PTC^Kdm8KO^*, n=10) and liver (right, *PTC^NT^*, n=13; *PTC^Kdm8KO^*, n=9) metastases in each *PTC* mouse of the indicated group. (**I**,**J**) Representative immunofluorescent images (top, **I**) with select regions (bottom, **I**) and the quantification of Kdm8 and Galectin-4 in 168,567 Tomato+ segments (**J**). Dotted line demarcates differentiated (Galectin-4+) region (bottom, **I**). The result is representative of tumors from n = 3 *PTC^Kdm8KO^* mice. (**K**) Representative H&E and immunohistochemistry (IHC) of Kdm8, and Vimentin of serial sections from a liver (top, **K**) and a lung (bottom, **K**) metastasis. (**L**) Zoomed-in images of the selected region of the liver metastasis highlighted by the red rectangle in K. (**M**,**N**) Representative H&E and IHC of Kdm8, Vimentin, and Tomato of serial sections from a primary PDA region (**M**) and the quantification of Kdm8 and Vimentin in 1,415 and 3,372 Tomato+ regions from 4 *PTC^NT^*(top) and 4 *PTC^Kdm8KO^* (bottom) mice, respectively (**N**). The Spearman correlation efficient ρ is shown. NS, not significant. Dotted lines demarcate metastases.

### CRISPR-mediated Kdm8 targeting induces a profound loss of differentiation and widespread metastatic lesions in autochthonous PDA

Compared to the *PTC^NT^* cohort, *PTC^Kdm8KO^*mice developed dramatically higher metastatic disease burden, including numerous micrometastases in the lungs (Figures 2F-2H and S2A-S2C). Of note, no mice in the *PTC^NT^* cohort developed any histologically discernable liver metastasis, consistent with previous reports that *Kras^G12D^*;*Trp53*-null PDA tumors are rarely metastatic (Figures 2G, 2H, and S2A) ^59,60^. Conversely, two-third of the *PTC^Kdm8KO^* mice had macroscopically evident metastases in the liver (*PTC^NT^*, n=0/13 vs *PTC^Kdm8KO^*, n=6/9, *p*=0.001 by two-sided Fisher’s exact test).

To gain further insights into the mechanism underlying this profound difference in metastasis, we examined the histological features of the primary PDA using Hematoxylin and Eosin staining (H&E), immunohistochemistry (IHC), and immunofluorescence (IF). While the *PTC^NT^* tumors exhibited homogeneous Kdm8 staining across histologically distinct regions, Kdm8 expression mostly restricted to fully differentiated glandular areas in the *PTC^Kdm8KO^*PDA (Figure S2E). Indeed, quantitative analysis revealed a strong positive correlation between the classical subtype marker Galectin-4 and Kdm8 (Figures 2I, 2J, and S2F). This suggests that *Kdm8* is required to maintain fully differentiated morphology in PDA, prompting us to test the hypothesis that the Kdm8-negative, less differentiated cancer cells might have transitioned into a mesenchymal state through EMT. To do this, we performed IHC across multiple primary and metastatic *PTC^Kdm8KO^* tumors. Indeed, *PTC^Kdm8KO^*metastases often lost Kdm8 expression while upregulating the EMT marker Vimentin (Figures 2K and 2L). Furthermore, quantification across thousands of histologic regions in primary tumors revealed a striking anti-correlation between Vimentin and Kdm8 IHC (Figures 2M, S2D, 2N). These results document a general enrichment of *Kdm8*-depleted malignant cells in the metastatic tumors of *PTC^Kdm8KO^* mice, suggesting enhanced metastatic fitness upon *Kdm8* loss. In summary, Kdm8 restrains phenotypic reprogramming by limiting the loss of differentiation and expression of the EMT marker Vimentin, thereby potently suppressing metastatic disease.

### The demethylase function of Kdm8 is critical for hypoxia-induced transcriptomic changes and metastatic progression

To examine the comprehensive gene expression change induced by *Kdm8* depletion, we comparatively profiled the full transcriptomics of control and sh*Kdm8* 688M cells cultured in normoxia (ambient O_2_) and hypoxia (0.5% O_2_) by performing bulk RNA-Seq analysis. Consistent with the IHC results, *Kdm8* knockdown in normoxia led to an upregulation of several hundred genes significantly enriched in EMT (Figures 3A and S3A). In contrast, *Kdm8* knockdown significantly decreased the classical-like signature ^38^, aligning with the loss of the differentiated state ^61,62^ (Figures 3A, 2I and 2J). Of note, *Kdm8* or *KDM8* knockdown in murine and human PDA cell lines, respectively, reduced the expression of the rate-limiting enzyme *Nsdhl* or *NSDHL* for cholesterol biosynthesis ^63^, which may explain the loss of glandular morphology in *Kdm8*-depleted PDA cells (Figures S3B). Interestingly, *Kdm8* knockdown in normoxia largely recapitulated the hypoxia-induced gene expression program, suggesting that a significant aspect of the hypoxic response in PDA cells is likely mediated through the suppression of Kdm8 function (Figures 3A, 3B, and S3C). Consistent with this notion, *Kdm8* knockdown in hypoxia failed to induce any Geneset Enrichment Analysis (GSEA) Hallmark transcriptomic programs, presumably due to the already diminished Kdm8 demethylase activity under hypoxic conditions (Figures 3A and S3D).

**Fig. 3.**
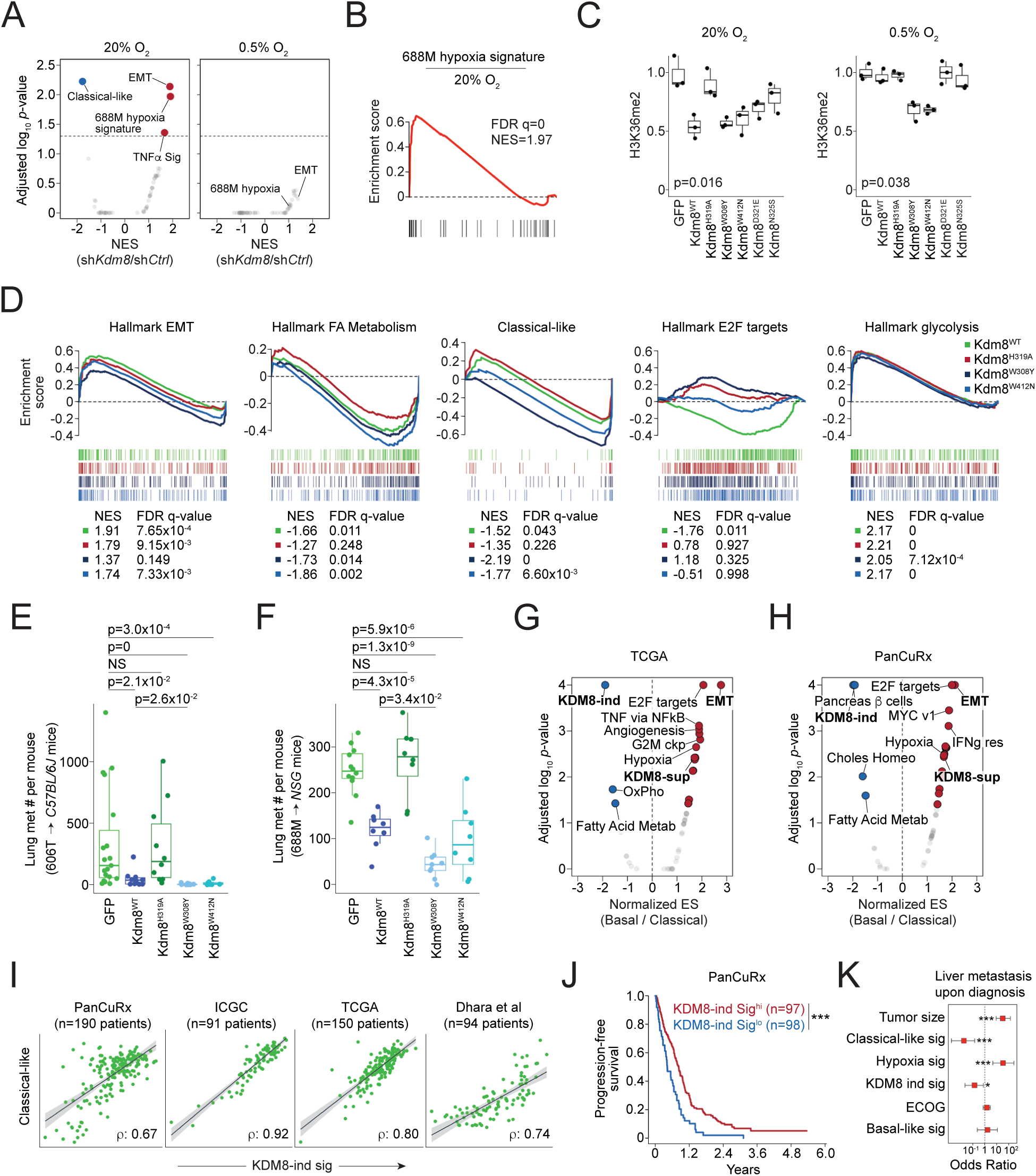
Kdm8 demethylase function is critical to hypoxia-induced transcriptomic reprogramming and metastatic progression. (**A**,**B**) Gene set enrichment analysis (GSEA) of the classical, basal, 688M hypoxia gene signature (**B**), and the hallmark GSEA signatures enriched in *Kdm8* knockdown 688M cells (compared to sh*Control*) cultured in normoxia (left, **A**) or hypoxia (right, **A**). Normalized enrichment score (NES) for the gene signatures and the corresponding FDR-adjusted p-values are shown. sh*Ctrl*, control sh*RNA*. Dotted line represents adjusted p=0.05. (**C**) Densitometric quantification of normalized H3K36me2 immunoblots (in *Kdm8* knockout (*Kdm8KO*) 688M cells upon the re-expression of GFP (as control) and the indicated Kdm8 variants in normoxia (left) or hypoxia (right). Results from three repeated experiments are shown. (**D**) GSEA of the indicated hallmark signatures enriched in hypoxic (compared to normoxic) *Kdm8KO* 688M cells expressing Kdm8^WT^, Kdm8^H319A^, Kdm8^W308Y^, and Kdm8^W412N^. NES for the gene signatures and the corresponding FDR-adjusted q-values are shown. (**E**,**F**) Counts of lung metastases in each *NSG* mouse orthotopically transplanted with *Kdm8KO* 606T (**E**, GFP, n=19; Kdm8^WT^, n=10; Kdm8^H319A^, n=10; Kdm8^W308Y^, n=10; Kdm8^W412N^, n=10) or 688M (**F**, GFP, n=12; Kdm8^WT^, n=8; Kdm8^H319A^, n=8; Kdm8^W308Y^, n=9; Kdm8^W412N^, n=8) murine PDA cells expressing the indicated Kdm8 variants or GFP as control. Kruskal-Wallis test (**E**, p=5.6×10-7) and one-way ANOVA (**F**, p=6.1×10-11) were used followed by Dunn’s and Dunnett’s test for multiple comparisons, respectively. (**G**,**H**) GSEA of the KDM8-induced (KDM8-ind) and KDM8-suppressed (KDM8-sup) gene signatures, and the hallmark GSEA signatures as in (**A**) enriched in bulk tumors defined as basal in comparison with those of the classical subtype from the TCGA (**G**) and PanCuRx (**H**) datasets. (**I**) Relationships between the KDM8-induced gene signature score (KDM8-ind sig) and classical subtype score across the indicated PDA bulk tumor RNA-Seq datasets. The Pearson correlation efficient ρ of each dataset is shown. (**J**) Kaplan-Meier survival analysis of the KDM8-induced gene signature score high (hi, defined as those higher than the IQR Q2 value) and low (lo) PDA patients from the PanCuRx data set. p=9.5×10-5 by log rank test. (**K**) Associations between the status of liver metastasis at diagnosis (dependent variable) and tumor size, classical and basal subtype gene signatures, hallmark hypoxia, KDM8-ind sig, or Eastern Cooperative Oncology Group performance status (ECOG) in the PanCuRx cohort (n=195). *, p<0.05; ***, p<0.001 by univariate logistic regression. Dotted line represents odds ratio=1.

To further test this hypothesis, we created Kdm8 hypermorphs by overlaying the JmjC domains of human KDM8 with two other KDM family members known for their high O_2_ binding affinities (KDM4A and KDM6B) ^64–66^. We identified four non-conserved residues, Y^177/1379^ (KDM4A/KDM6B) -> W^310^ (KDM8), E^190/1392^ (KDM4A/KDM6B) -> D^323^ (KDM8), S^196/1398^ (KDM4A/KDM6B) -> N^327^ (KDM8), and N^290/1484^ (KDM4A/KDM6B) -> W^414^ (KDM8), which engage either α-ketoglutarate or Fe^+2^ according to previous reports and may enhance KDM8 demethylase function upon replacement ^67,68^ (Figure S3E). Compared to the wildtype counterpart (Kdm8^WT^), which is functional only in normoxia, expression of Kdm8^W308Y^ or Kdm8^W412N^ (murine orthologs to human KDM8^W310Y^ and KDM8^W414N^, respectively) in the *Kdm8* knockout 688M cells reduced H3K36me2 level in both normoxia and hypoxia, indicating heightened demethylase activities (Figures 3C, S3F, and S3G). In support of this, *Kdm8*-deficient 688M cells expressing either Kdm8 hypermorph demonstrated reduced sensitivity to hypoxia-induced cell cycle arrest compared to the Kdm8^WT^-expressing counterparts (Figure S3H). Additionally, expression of the hypermorphic Kdm8 variants abolished hypoxia-induced changes in multiple transcriptional programs, including the upregulation of EMT and downregulation of cell cycle (Figure 3D). In contrast, hypoxia-induced suppression of the classical-like program and fatty acid metabolism was sensitive to the expression of the demethylase inactive variant Kdm8^H319A^ (Figure 3D). Consistent with the major role of the HIF pathway in facilitating anaerobic glycolysis, hypoxia-induced glycolysis appeared to be unaltered by changes in Kdm8 activity (Figure 3D) ^69^.

We next examined the changes upon expressing these Kdm8 variants in the context of metastasis. To see whether the hypermorphic Kdm8 variants have an even stronger anti-metastatic effect, we performed orthotopic tumor studies using *Kdm8* knockout 688M and 606T PDA cells. In both models, expression of Kdm8^H319A^ did not alter the level of lung metastasis contrasted by the Kdm8^WT^ counterpart, supporting the role of the Kdm8 JmjC domain in restoring a low metastatic state (Figures 3E, 3F, S3I, and S3J). Notably, expression of the hypermorphic Kdm8^W308Y^ variant led to significantly reduced lung metastatic burdens compared to the Kdm8^WT^ counterparts in both models (Figures 3E and 3F). In summary, these observations suggest that hypoxia promotes metastasis through the suppression of Kdm8, and that Kdm8 activity is a major rheostat that controls PDA metastatic ability.

### KDM8-induced gene signature predicts survival and metastatic disease in human PDA

Our results indicate that Kdm8 demethylase function suppresses metastasis and maintains a differentiation in PDA in murine models. We next sought to determine whether the KDM8-induced gene signature predicts survival and disease behavior in human PDA. *KDM8* is relatively homogeneously expressed, thus we performed correlative analyses using KDM8-regulated gene signatures. Comparative profiling of bulk RNA-Seq data from The Cancer Genome Atlas (TCGA) ^70^ and the Ontario Institute for Cancer Research (PanCuRx Translational Research Initiative) ^38^ reveals a consistent enrichment of the KDM8-induced gene signature in the classical subtype (Figures 3G and 3H). Indeed, classical-like and KDM8-induced signatures were well correlated across four different PDA cohorts, despite minimum overlap between the two gene lists ^71,72^ (Figures 3I and S3K). Using single nuclei transcriptomes isolated from a separate PDA cohort ^73^, we found a striking correlation between the classical-like and KDM8-induced gene signatures in malignant cells, thus eliminating the possibility that the correlations in the bulk RNA-Seq datasets were driven by non-malignant cell types in tumors (Figures S3L and S3M). In the PanCuRx cohort, the KDM8-induced gene signature is among the top predictors for prognosis, with high scores correlating with better outcomes (Figure 3J). Importantly, patients with a high KDM8-induced gene signature or classical-like score have significantly reduced odds of liver metastasis at diagnosis (Figure 3K). As anticipated, the hypoxia signature is a potent risk factor for liver metastasis ^4^ (Figure 3K). These results suggest that in human PDA, KDM8 function maintains the classical program and is associated with reduced metastasis prior to therapeutic intervention.

### Malignant state plasticity is regulated by Kdm8 in vivo

In PDA, transcriptomic programs define distinct disease subtypes that are clinically relevant ^27,38,74,75^. The basal-like subtype (*a.k.a.* squamous-like or quasi-mesenchymal) is associated with advanced stage ^38^, resistance to treatment ^73,76–78^, and serves as an independent predictor for poor prognosis in multiple studies ^4,27,74,75,79^. More recently, the neural-like progenitor malignant cell program has been identified as one of the most resistant subtypes in chemotherapy and radiotherapy, correlating with poor outcomes in independent cohorts ^73^. Despite this, the role of the hypoxia-Kdm8 axis in the plasticity of malignant programs remains undefined.

To investigate the transcriptomic state plasticity in the *PTC^Kdm8KO^*mice, we performed single cell RNA-Seq (scRNA-Seq) on *ex vivo* malignant cells isolated using fluorescence-activated cell sorting (FACS) for Tomato-positive and lineage marker (Cd45, Ter119, F4/80, Cd31) negative population (Figures 4A and S4A) ^11^. The scRNA-Seq dataset comprises 32,878 cells with a median of 17,107 unique molecular identifiers (UMIs) and 4,061 genes detected across 11 tumors and metastatic lesions (5 primary PDA tumors, 3 disseminated cancer cell samples from the ascites, and 3 lung metastases). Utilizing previously reported signatures, we uncovered 9 malignant programs that closely resembled human PDA, including classical-like, EMT, classical/basal hybrid, basal/hypoxia hybrid, hypoxia, Myc, neuroendocrine, and transitional states (Figures 4B and S4B) ^73^. Furthermore, by employing morphologically associated gene signatures that define distinct PDA histological patterns ^61^, we confirmed that the classical, classical/basal hybrid, and transitional states predominantly resembled morpho-biotype A (ductal or glandular-like structure), while the EMT state aligned with morpho-biotype B (ill-defined glands, Figure S4C).

**Fig. 4.**
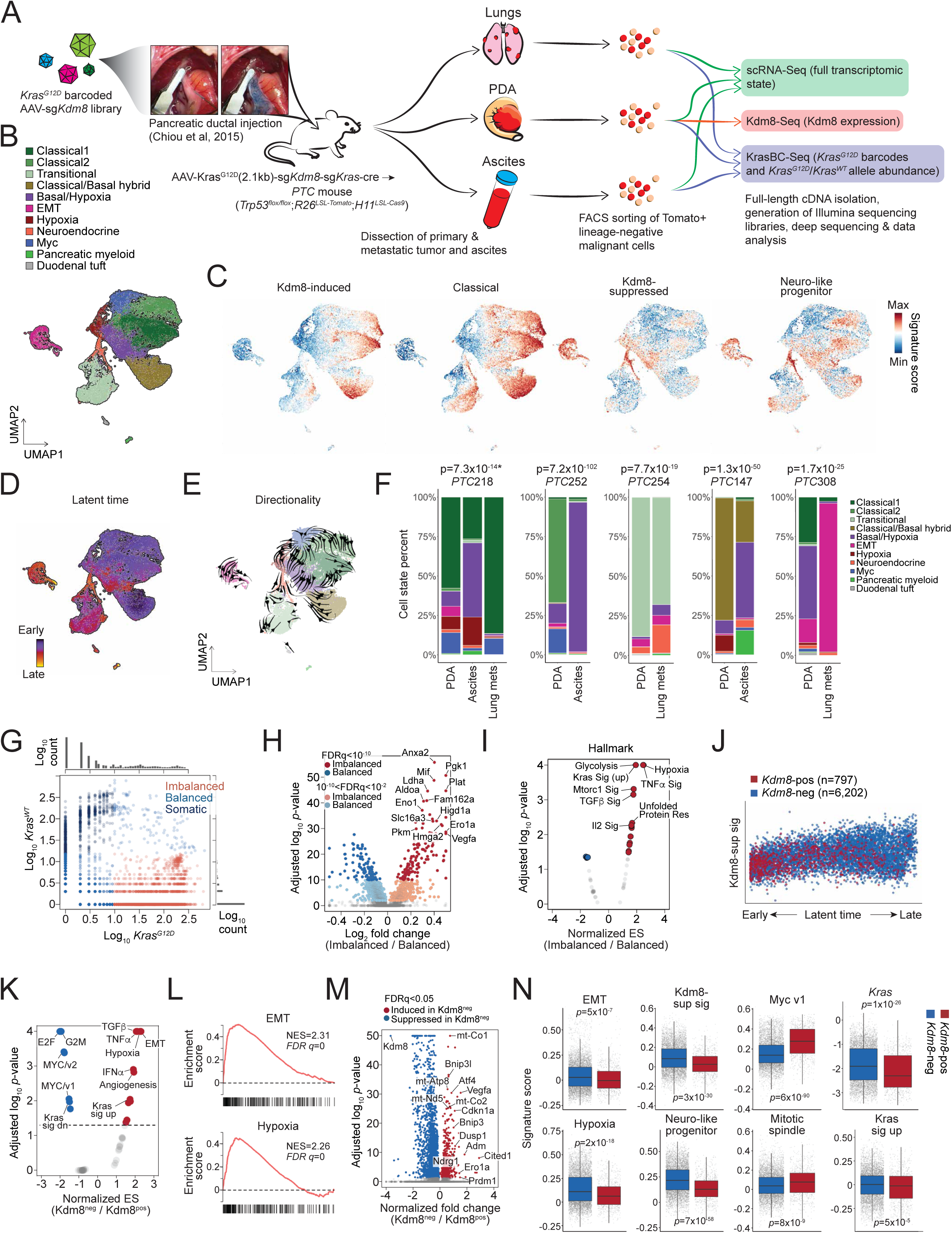
Malignant cell state plasticity is regulated by Kdm8 *in vivo*. (**A**) Schematic of the single-cell sequencing experiment using dissociated malignant cells from the *PTC^Kdm8KO^*mice. (**B**-**E**) UMAP embeddings of single-cell profiles of all Tomato+ malignant cells derived from 11 tumors and metastatic lesions colored by post hoc cell cluster annotations using Seurat 4.1.3 (**B**), indicated malignant transcriptomic programs (**C**), and latent time (**D**) or pseudotime trajectory (**E**). (**F**) Distributions of the identified malignant programs as in (**B**) across the 11 tumors and metastatic lesions. Stacked bar plots showing the proportions (Y-axis) of malignant programs across samples isolated from 5 *PTC^Kdm8KO^* mice. Fisher’s exact test p-values for the association between the advanced malignant state (EMT, neuroendocrine, and basal/hypoxia) and metastasis are shown (top). *, association between the malignant state and ascites is shown. (**G**) Log *Kras^G12D^* and *Kras^WT^* transcript abundance in Tomato+ malignant cells isolated from *PTC^Kdm8KO^* tumors (n=7,194 cells) that are colored according to their *Kras^G12D^*/ *Kras^WT^* allelic conformation. (**H**,**I**) Differentially expressed genes (H) and GSEA for the hallmark gene signatures enriched in malignant cells with *Kras^G12D^*/ *Kras^WT^*allelic imbalance compared to counterparts without the allelic imbalance (**I**). Somatic cells shown in Fig. **G** were excluded from this analysis. ES, enrichment score. (**J**) Increase in Kdm8-suppressed gene signature score along the latent time trajectory defined in (**D**). Cells are colored by the *Kdm8* status using Kdm8-Seq. Kdm8-sup sig, Kdm8-suppressed gene signature. (**K**,**L**) GSEA of the hallmark gene signatures independently identified transcriptomic programs, including EMT and hypoxia (**L**), that are enriched in Kdm8 negative (*Kdm8^neg^*) malignant cells (**K**). (**M**) Differentially expressed genes identified by comparing *Kdm8^neg^* over *Kdm8^pos^* cancer cells. (**N**) Abundance of select malignant programs (Hwang et al, 2022 Nat Genet), GSEA hallmark gene signatures, *Kras* expression, and Kdm8-suppressed gene signature in *Kdm8* positive and negative cancer cells.

Consistent with human PDA, a striking positive correlation between the states defined by KDM8-induced gene signature and the classical-like signature was observed (Figures 4C and S4D). Surprisingly, the expression of KDM8-suppressed genes showed a high correlation with the neuro-like progenitor signature ^73^, even though the two gene lists minimally overlapped, implicating the role of hypoxia in suppressing Kdm8 and the subsequent induction of this aggressive malignant cell program (Figures 4C, S4D, and S3K). Of note, a low Myc score, which corresponds to a low proliferative state, coincided with a high abundance of genes suppressed by Kdm8, suggesting a cell cycle-antagonizing effect following Kdm8 inactivation (Figure S4D). We further validated this by demonstrating reduced cell proliferation upon *Kdm8* knockdown in multiple PDA cell lines (Figures S4E).

Finally, we examined differentiation trajectories across the malignant cell states by performing RNA velocity analysis to determine inferred latent time and differentiation directionality (Figures 4D and 4E) ^80,81^. Consistent with previous findings ^6,82^, our data revealed a transcriptomic continuum spanning across the Myc (near root), classical-like, and classical/basal hybrid states (early) through to hypoxia and transitional states (late), and ultimately the terminal states, which include basal/hypoxia hybrid, neuroendocrine, and EMT programs (Figures 4D, 4E, and S4F). Along the latent time trajectory, EMT, neuro-like progenitor, and Kdm8-suppressed gene expression programs displayed rising patterns, while classical-like and Myc gene signatures declined (Figure S4F). As anticipated, disseminated and metastatic lesions show significant enrichment of the terminal programs compared to their autologous primary PDA (Figure 4F). These findings support a general model of PDA plasticity in which the Myc and classical-like programs represent early states that progress into late (hypoxia and transitional) and terminal states (basal/hypoxia hybrid, neuroendocrine, and EMT).

In addition to scRNA-Seq for cell state profiling, the full-length 10X cDNA libraries were subjected to paired analysis measuring the abundance of the barcoded *Kras^G12D^* alleles (KrasBC-Seq, Figures 4A and S4G). This paired measurement of barcoded *Kras^G12D^* and wildtype *Kras* alleles narrowed down the original scRNA-Seq dataset to 7,194 cells with at least one detected read count. As expected, the overall *Kras* transcript abundance quantified with KrasBC-Seq correlated with those in scRNA-Seq, albeit with much higher resolution (left, Figure S4H). Consistent with a previous report ^58^, we found that a vast majority of the metastatic lesions derived from the largest primary PDA clonotypes. Multiple malignant programs were detected within individual *Kras^G12D^* clonotypes, including both classical- and basal-like subtypes, highlighting the importance of the tumor microenvironment over the cell of origin in driving transcriptomic plasticity (Figure S4I). Given that *KRAS* allelic imbalance correlates with PDA progression and is linked to advanced PDA subtypes and metastasis ^38,39,83^, we sought to examine the level of *Kras* allelic imbalance relative to distinct malignant cell states in the *PTC^Kdm8KO^* mice. Leveraging the barcode strategy that enabled unambiguous calling of *Kras^G12D^* and wildtype *Kras* alleles, we clearly distinguished malignant cells with balanced versus imbalanced *Kras^G12D^*/*Kras^WT^* expression (Figure 4G). A small fraction (n= 621/7,194) of the cells exhibited higher *Kras^WT^* expression compared to the barcoded *Kras^G12D^*alleles, presumably comprised primarily nonmalignant cells in the current PDA model. Interestingly, comparative profiling using GSEA Hallmark signatures uncovered hypoxia and Kras signaling as the top enriched programs in malignant cells with *Kras* allelic imbalance (Figures 4H and 4I). Furthermore, *Kras* allelic imbalance was consistently enriched in hypoxic cells across all *PTC^Kdm8KO^* mice, indicating that hypoxia might exert strong selective pressure favoring increased *Kras^G12D^*/*Kras^WT^*ratios (Figure S4J).

To directly assess the impact of *Kdm8* depletion on transcriptomic state plasticity, we performed paired *Kdm8* targeted sequencing with the scRNA-Seq cDNA libraries (Kdm8-Seq, Figure 4A). Consistent with the low Kdm8 expression by IHC observed in metastases (Figures 2K and 2L), metastatic lesions yielded minimal PCR amplicon with poor library quality (data not shown). The paired analysis successfully generated targeted *Kdm8* sequencing libraries from a total of 5 primary *PTC^Kdm8KO^* PDA tumors that also had scRNA-Seq and KrasBC-Seq datasets available (n=6,999 cells). Not surprisingly, we noted an exceedingly high correlation between *Kdm8* measured with Kdm8-Seq and scRNA-Seq (right, Figure S4H) and a bimodal distribution of Kdm8 abundance with 11.4% (n=797/6,999) of the cells expressing *Kdm8*, comparable to results obtained using the Surveyor nuclease assay (Figure S4K). Strikingly, Kdm8^pos^ cancer cells were unevenly scattered along the latent time trajectory, with most residing near the root and decreasing in density towards late and terminal stages, suggesting that *Kdm8* loss drove cancer cells into advanced transcriptomic states (Figure 4J). Indeed, comparison of Kdm8^neg^ versus Kdm8^pos^ cells using GSEA hallmark signatures and differential gene analysis revealed significantly enriched hypoxia, EMT, and Kras-upregulated genes, while E2F targets, G2M checkpoint, and Kras-downregulated gene signatures were depleted (Figures 4K-4N and S4L). These results provided evidence supporting the *in vivo* role of *Kdm8* in suppressing advanced malignant programs and Kras signaling.

### Kdm8 depletion recapitulates hypoxia-induced epigenetic reprogramming

To define the global epigenetic changes resulted from hypoxia-induced Kdm8 suppression, we performed chromatin immunoprecipitation sequencing (ChIP-Seq) to quantify various histone modifications, including H3K36me2 and H3K27 trimethylation (H3K27me3), which are likely impacted by Kdm8 inactivation based on previous findings ^49–51^. We quantitatively measured H3K36me2 and H3K27me3 global changes in control and sh*Kdm8* 688M cells cultured in normoxia and hypoxia. In the same experiment, we also performed H3K27 acetylation (H3K27ac) ChIP-Seq and assay for transposase-accessible chromatin with sequencing (ATAC-Seq) to simultaneously assess these chromatin features as potential secondary changes that might explain transcriptomic reprogramming following *Kdm8* knockdown or hypoxia. *Kdm8* knockdown induced a marked increase in H3K36me2 levels across 7,952 and a decrease in 7,140 genomic regions (Figure 5A). Similarly, global changes in H3K27me3 were observed upon *Kdm8* knockdown (Figure 5A). Of note, short-term exposure to hypoxia resulted in substantial global changes in both H3K36me2 and H3K27me3 (Figure S5A). In addition, the global changes in H3K27ac upon *Kdm8* knockdown were only partially mirrored by hypoxia (Figures 5A and S5A). Finally, *Kdm8* depletion has a minimal impact on global chromatin accessibility, whereas hypoxia induced robust genome-wide changes (Figures 5A and S5A).

**Fig. 5.**
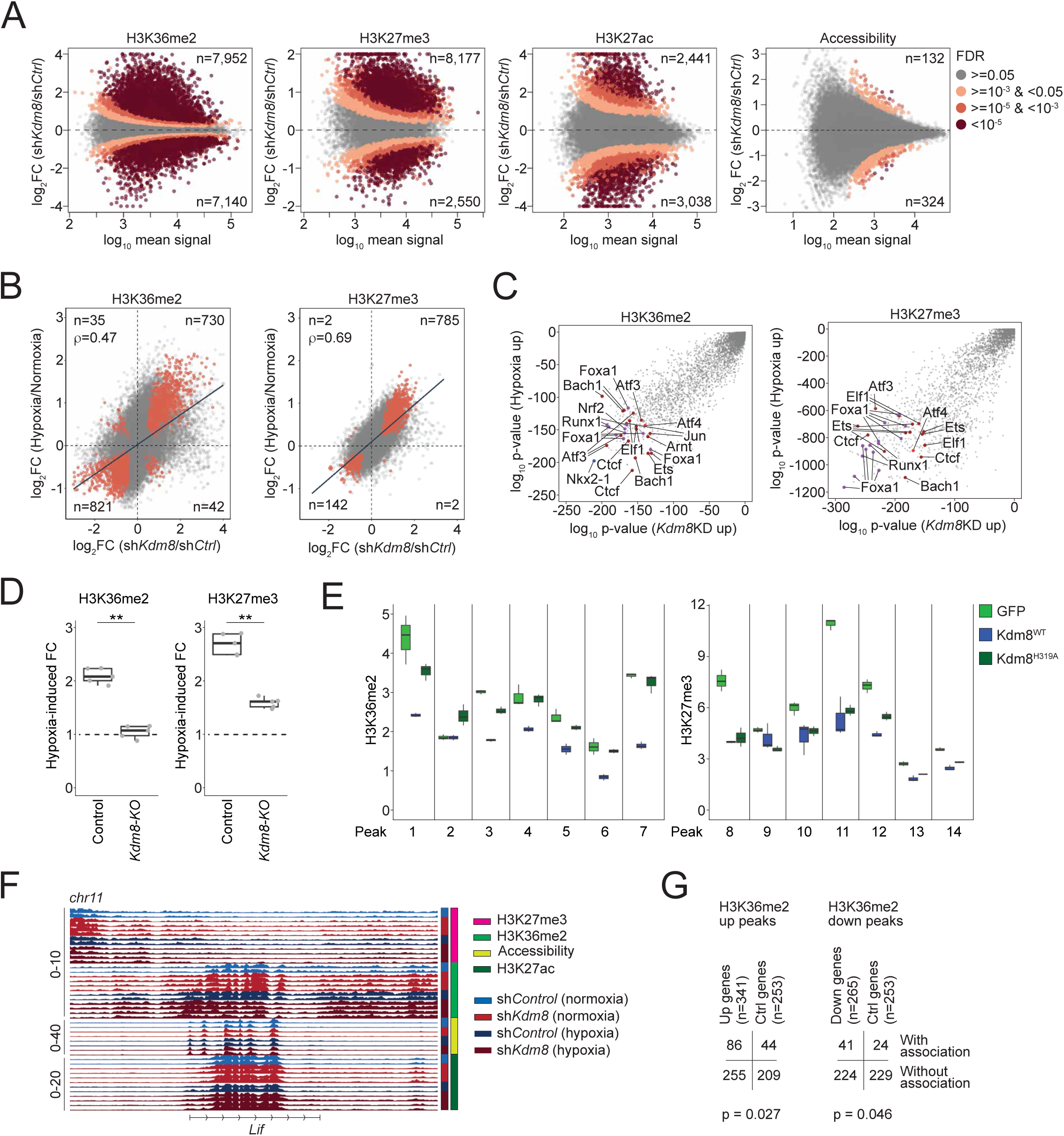
*Kdm8* depletion recapitulates hypoxia-induced epigenetic reprogramming. (**A**) *Kdm8* knockdown (*Kdm8KD*)-induced differential H3K36me2, H3K27me3, and H3K27ac ChIP-Seq signals, as well as chromatin accessibility measured by ATAC-Seq (left to right), plotted against the log mean read counts per region (x-axis). Dotted lines represent zero log_2_ fold change (FC), with positive values representing *Kdm8KD*-induced increases. FDR, false discovery rate that a differential region reveals significant change following *Kdm8* knockdown. The number of significantly altered regions (FDR < 0.05) is indicated at the top and bottom of each plot. (**B**) Relationships between *Kdm8KD*-induced (x-axis) and hypoxia-induced (y-axis) fold changes for H3K36me2 (left) and H3K27me3 (right). Regions that are significantly altered by *Kdm8KD* and hypoxia are highlighted and the numbers in each quadrant are shown. Pearson correlation coefficients (ρ) are shown. (**C**) Homer transcription factor (TF) motif enrichment analysis showing the probability of TF motif enrichment in *Kdm8KD*-induced (x-axis) and hypoxia-induced (y-axis) regions for H3K36me2 (left) and H3K27me3 (right). TF motifs associated with high-altitude adaptation (Xin et al, 2020 Nat Commun), stress response, and PDA progression are highlighted. (**D**) Densitometry of immunoblots quantifying hypoxia-induced fold change (FC) of H3K36me2/H3 (left) and H3K27me3/H3 (right) ratios in control (parental) and *Kdm8* knockout (*Kdm8*-KO) 688M cells. (**E**) Quantitative PCR analysis of H3K36me2 (left) and H3K27me3 (right) ChIP signals in *Kdm8*-KO 688M cells expressing the indicated Kdm8 variants, measured across genomic regions (Peaks 1 to 14) prioritized by ChIP-seq. (**F**) H3K36me2, H3K27me3, H3K27ac, and ATAC-Seq signals in control and *Kdm8KD* 688M cells cultured in normoxia or hypoxia at the *Lif* locus. (**G**) Contingency tables showing the number of genes co-induced by *Kdm8KD* and hypoxia that overlap with regions induced by either hypoxia or *Kdm8KD* (left), and genes co-suppressed by *Kdm8KD* and hypoxia that overlap with regions suppressed by either condition (right). Control (*Ctrl*) genes are defined as those with p > 0.9 for both hypoxia and *Kdm8KD*-induced changes.

To gain further insights, we examined the distribution of regions with changes in H3K36me2 and H3K27me3. Interestingly, hypoxia suppressed 27.4% of all H3K36me2 regions and 10.1% of all H3K27me3 regions that are at least 1 kb away from the nearest transcription start site (TSS) or gene distal, which was more than double the suppression observed in their gene-proximal counterparts (bottom, Figure S5B; top, Figure S5C). Similar patterns of the reduced H3K36me2 and H3K27me3 regions were observed following *Kdm8* knockdown (top, Figure S5B; top, Figure S5C). Notably, suppression of these H3K36me2 gene-distal regions may be a secondary effect of *Kdm8* knockdown and could be attributed to increased *Kdm2a* expression following *Kdm8* depletion or hypoxia (not shown), consistent with a prior report of Kdm2a’s demethylase activity at gene-distal H3K36me2 sites ^84^. In contrast, an opposite pattern was observed for hypoxia-promoted H3K36me2 and H3K27me3 regions, as well as for the differential H3K27me3 regions induced by *Kdm8* knockdown, with these regions being predominantly gene-proximal rather than gene-distal (Figure S5B; bottom, Figure S5C). Remarkably, a striking correlation between the fold changes of all regions induced by hypoxia and *Kdm8* knockdown was noted, indicating that *Kdm8* knockdown itself is sufficient to recapitulate most hypoxia-induced changes in these histone marks (Figure 5B). Furthermore, assessment of enriched transcription factor motifs within H3K36me2 and H3K27me3 regions induced by hypoxia or *Kdm8* knockdown revealed co-enrichment of multiple binding sites for factors critical in high-altitude adaptation, stress response, and PDA progression (Figure 5C) ^13,85–87^. Consistent with Kdm8’s function in hypoxia-induced epigenetic changes, *Kdm8* depletion through CRISPR knockout completely and partially abolished hypoxia-induced accumulation of H3K36me2 and H3K27me3, respectively (Figures 5D and S5D). We validated the role of Kdm8 JmjC domain function in modulating H3K36me2, but not H3K27me3, in multiple Kdm8-regulated genomic regions by reintroducing the WT or enzymatically inactive Kdm8^H319A^ variant in *Kdm8*-deficient cells (Figures 5E). In summary, the impact of *Kdm8* inactivation mirror a large part of hypoxia-induced chromatin changes, strongly supporting the model of the hypoxia-Kdm8 axis in epigenetic reprogramming.

To gain additional insights into the regulation of gene expression during the hypoxic response, we focused on differential gene proximal regions induced by *Kdm8* depletion and hypoxia (Figure S5B). We identified numerous genes, including *Lif*, where the expression changes induced by hypoxia or *Kdm8* knockdown aligned with changes in H3K36me2 (Figures S3A and 5F). Consistent with H3K36me2’s role in actively transcribed loci, both hypoxia and *Kdm8* knockdown resulted in increased *Lif* expression, coinciding with elevated H3K36me2 signals at multiple peaks near the *Lif* locus (Figures S3A and 5F). Overall, 86 of the 341 (25.2%) genes that were induced by both hypoxia and *Kdm8* knockdown were associated with hypoxia- or *Kdm8* knockdown-induced H3K36me2 proximal regions, whereas 41 of the 265 (15.5%) suppressed genes were linked to hypoxia- or *Kdm8* knockdown-repressed H3K36me2 proximal regions (Figure 5G). Comparative profiling using GSEA c2 curated gene sets uncovered an H3K27me3-regulated gene signature ^88^ as the top depleted program upon *Kdm8* knockdown in normoxia, suggesting that the hypoxia-Kdm8 axis might regulate the transcriptomes by orchestrating different epigenetic changes at different genomic sites (Figure S5E). Consistent with this, we noted a coordinated increase in H3K27ac and a decrease in H3K27me3 within the H3K36me2 regions induced by both *Kdm8* knockdown and hypoxia (n=3,973 regions). Conversely, the suppressed H3K36me2 regions (n=3,838) were associated with reduced H3K27ac and enhanced H3K27me3 signals (Figures S5F and S5G). Collectively, these results support a mechanism by which the hypoxia-Kdm8 axis induces major transcriptomic reprogramming by coordinating global epigenetic changes.

### The hypoxia-Kdm8 axis promotes Kras copy-number gain through accelerated CIN

CIN is a well-established driver of metastasis in multiple solid tumor models ^89^. Given the critical role of *jmjd-5* in preserving chromosome integrity in *C. elegans* ^55^ and the association of *JMJD5* germline mutations with DNA replication stress in humans ^54^, we hypothesize that hypoxia promotes CIN by suppressing Kdm8 demethylase activity. To assess the role of Kdm8 in regulating CIN, we applied complementary assays: (i) WGS to measure focal CNA and aneuploidy, (ii) DAPI staining of genomic DNA to quantify micronuclei formation, and (iii) immunofluorescence to analyze mitotic spindle organization during anaphase.

We first assessed micronuclei formation in PDA cells following *KDM8* knockdown. In both murine and human PDA cell lines, knockdown of *Kdm8* or *KDM8*, respectively, increased micronuclei under normoxic conditions (Figure 6A). In 688M PDA cells, *Kdm8* knockdown alone recapitulated hypoxia-induced preponderance of micronuclei, indicating that hypoxia may promote CIN through Kdm8 suppression (Figure 6A). Whole chromosome painting in 688M cells revealed a tetraploid karyotype with increased double minutes derived from chromosome 6, whereas chromosome 11 remained unaffected, indicating that *Kdm8* knockdown drives chromosome-specific structural alterations induced by the hypoxia-Kdm8 axis (Figures 6B and 6C). Sporadic chromothripsis events were observed in *Kdm8* knockdown cells, absent controls, albeit without statistical significance (data not shown). These results indicate that *Kdm8* inactivation promotes CIN in PDA cells.

**Fig. 6.**
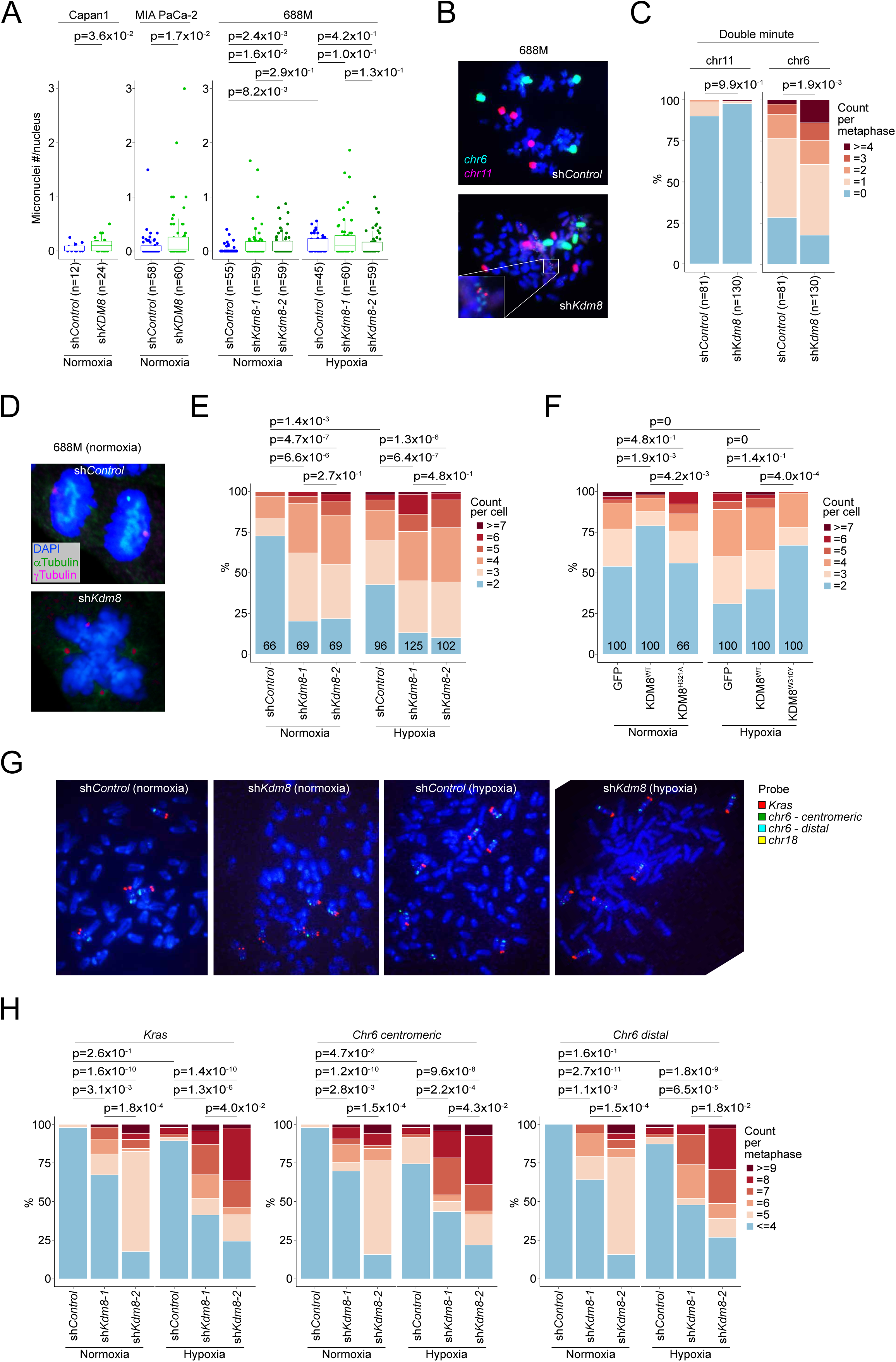
The hypoxia-Kdm8 axis promotes *Kras* copy-number gain through accelerated chromosomal instability. (**A**) Quantification of micronuclei formation, shown as normalized ratios of micronuclei counts per nucleus, in human PDA cells (Capan-1 and MIA PaCa-2; *control* vs. sh*KDM8*-expressing) cultured in normoxia, and murine 688M PDA cells (*control* vs. sh*Kdm8*-expressing) cultured in normoxia or hypoxia. (**B**,**C**) Representative images (**B**) and quantifications (**C**) of whole chromosome 6 and 11 painting in control and sh*Kdm8*-expressing 688M cells. The proportions of metaphase cells in control and the sh*Kdm8* group associated with the indicated numbers of chromosome 6- and 11-derived double minutes are shown, with p-values calculated using one-sided Wilcoxon signed-rank test on double minute tallies (**C**). (**D**,**E**) Representative images of anaphase cells in normoxia (**D**) and the proportions of control and *Kdm8* deficient 688M PDA cells cultured in normoxia or hypoxia with the indicated anaphase polarities quantified using γ-Tubulin immunofluorescence (IF, **E**). (**F**) Proportions of *KDM8* knockout Capan-1 cultured in normoxia (left) or hypoxia (right) with anaphase polarities quantified by Pericentrin IF expressing the indicated KDM8 variants. (**G**,**H**) Representative images (**G**) and quantifications (**H**) of fluorescent in-situ hybridization (FISH) using probes for Kras (red), chromosome 6 centromeric (green) and distal (cyan) regions, and chromosome 18 (yellow) in control (normoxia, n=49; hypoxia, n=50) and sh*Kdm8*-expressing 688M cells cultured in normoxia (sh*Kdm8-1*, n=53; sh*Kdm8-2*, n=51) or hypoxia (sh*Kdm8-1*, n=51; sh*Kdm8-2*, n=50). The proportions of control and the sh*Kdm8* metaphase 688M cells cultured in normoxia or hypoxia with the indicated FISH probe tallies are shown (**H**). Numbers in the plot indicate the number of cells quantified in each group (**E**,**F**). Unless otherwise stated, p-values were derived using two-sided Wilcoxon test for comparisons of fluorescent quantification between two groups or a Kruskal-Wallis test for comparisons among three or more groups, followed by Dunn’s test for sub-comparisons with Benjamini-Hochberg adjustment.

Next, to determine whether hypoxia compromises mitosis through the suppression of Kdm8 demethylase activity, we quantified spindle organization defects by immunofluorescence. Consistent with previous reports ^56,90^, *Kdm8* knockdown in both 688M and 606T murine PDA cell lines increased mitotic abnormalities, specifically multipolar spindle formation, a known cause of chromosome mis-segregation ^91^ (Figures 6D, 6E, and S6A). Similar spindle defects were observed in human Capan-1 cells and non-malignant MCF10A cells upon *KDM8* depletion (Figure S6A). Hypoxia promoted multipolar spindles during anaphase in control 688M cells, while *Kdm8* knockdown under normoxia produced the same phenotype, concordant with the micronuclei observations (Figure 6E). Importantly, expression of KDM8^WT^, but not the demethylase-inactive KDM8^H321A^ variant, in *KDM8* knockout Capan-1 cells restored spindle organization in normoxia (Figure 6F). Similarly, expression of the demethylase-inactive Kdm8^H319A^ variant in the murine PDA 688M cells had minimum effect on spindle polarity (Figure S6B). In hypoxia, expression of the hypermorphic KDM8^W310Y^ variant in human PDA cells or the corresponding Kdm8^W308Y^ variant in murine PDA cells - but not their WT counterparts - significantly reduced spindle defects, demonstrating that KDM8 demethylase activity is essential for maintaining mitotic fidelity and limiting CIN (Figures 6F and S6B).

Given the increased mitotic errors, we performed bulk and single-cell WGS (scWGS) to further examine karyotypic heterogeneity upon *Kdm8* knockdown. Interestingly, *Kdm8* depletion had distinct impacts on the ploidies of different chromosomes, suggesting that CNAs in select chromosomes may serve as preferred genomic features for *in vitro* growth (Figures S6C and S6D). Notably, chromosome 6, which contains *Kras*, was recurrently amplified following *Kdm8* knockdown using different hairpins (Figures S6D and S6E). In line with these observations and the increased Kras signaling in *PTC^Kdm8KO^*cancer cells (Figures 4K, 4N, and S4L), we hypothesize that hypoxia promotes *Kras* amplification through the suppression of Kdm8 demethylase function.

In patients with advanced PDA, the KDM8-induced gene signature is associated with reduced *KRAS* allelic imbalance, a feature largely driven by oncogenic *KRAS* amplification ^39^ (Figure S6F). To test this hypothesis, we assessed the impact of *Kdm8* knockdown or hypoxia on *Kras* CNAs using fluorescent in situ hybridization (FISH). Consistent with the WGS data, *Kdm8* knockdown with two independent hairpins resulted in significant chromosome 6 copy-number gains, relative to chromosome 6 copies quantified using a centromeric probe (Figures 6G and 6H). In contrast, reference chromosome 18 exhibited minimal ploidy changes following *Kdm8* knockdown (Figures S6G and S6C). Hypoxia also increased chromosome 6 ploidy in control 688M cells, though to a lesser degree. Both hypoxia and *Kdm8* knockdown increased *Kras* copy number along with a distal reference region on chromosome 6, consistent with chromosome 6 polyploidy. These data indicate that *Kras* copy-number gain arises through chromosome 6 amplification (Figures 6G and 6H). In agreement with this observation, *Kdm8* and *KDM8* knockdown in multiple murine and human PDA cell lines significantly increased sensitivity to the KRAS^G12D^ inhibitor MRTX1133 (Figures S6H and S6I). In summary, the hypoxia-Kdm8 axis accelerates CIN, leading to chromosome 6 polyploidy and subsequent *Kras* amplification.

## Discussion

This work reveals a novel mechanism by which PDA malignant cells co-opt select elements of the hypoxic response that drive metastatic spread. Our data suggest that by suppressing KDM8 demethylase function, hypoxia unleashes an adaptive program characterized by reduced cell division, increased CIN, loss of differentiation, and enhanced EMT. As a hallmark of cancer ^92^, CIN, enabled by ongoing mitotic errors, generates a pool of malignant cells with diverse genomic and phenotypic traits – some of which are readily selected by and survive environmental constraint. In the context of hypoxia, this model explains the adaptability of genomically unstable malignant cells to cope with the metabolic challenges within the tumor microenvironment. In support of this, our results show that *Kdm8* suppression in cancer cells induces the acquisition of higher *Kras* copy numbers, a heritable trait that facilitates the removal of hypoxia-induced reactive oxygen species through increased glutathione biosynthesis ^93^ and is preserved in metastases ^38^. Furthermore, hypoxia-mediated Kdm8 suppression promotes a metastatic program by facilitating EMT and loss of differentiation through (i) H3K36me2 upregulation and (ii) induction of CIN, both of which have been previously reported to reinforce an EMT-like phenotype ^84,89^. Restoration of the hypoxia-resistant, hypermorphic Kdm8 variant almost completely blocked metastasis and could provide a viable therapeutic strategy to reverse the highly metastatic behavior in PDA.

Consistent with previous reports ^49,50^, *Kdm8* depletion reduced the proliferation of PDA cells. In the *PTC* mice, *Kdm8* knockout in PDA cancer cells diminished the Myc gene signature, which was enriched near the root of the latent time trajectory. In human PDA, the MYC transcriptomic state shows strong negative correlations with the KDM8-suppressed gene signature, EMT, and hypoxic response (Figure S4D). This finding aligns with an earlier report analyzing bulk RNA-Seq data from 500 patients across more than two dozen carcinoma types, which revealed a strong negative correlation between EMT and proliferation signatures ^94^. Under normal circumstances, cells undergoing morphogenesis (e.g. EMT) are evolutionarily hardwired for cell cycle arrest to prevent severe developmental defects caused by mitosis checkpoint failure ^95,96^. Despite this, *MYC* is a PDA oncogene ^97^ that is amplified in metastases ^98,99^, which appears paradoxical given that suppressed MYC gene signature is part of the hypoxia-induced metastatic reprogramming. As a hallmark of cancer ^92^, unrestrained cell proliferation induces genomic instability in epithelial cancer cells undergoing EMT ^100^. Consistent with this, a mesenchymal state defined as Vimentin-positive is strictly required for CIN in PDA, irrespective of the genomic drivers used in the GEM models ^21^. Thus, in the context of hypoxia-induced EMT, it is conceivable that increased *MYC* dosage would be a highly favorable trait for promoting CIN and metastasis ^101,102^. Although beyond the scope of the study, it would be of interest to elucidate a potential mechanism by which EMT, regardless of its induction by hypoxia or other factors, fuels *MYC* amplification in promoting PDA metastasis.

High prevalence of *KRAS* and *TRP53* genetic mutations in human PDA underscores the development of PDA GEM models ^103^. Due to their similarity to human disease, *KPC* mice continue to play a pivotal role in advancing our understanding of PDA biology. However, these GEM models are labor-intensive and highly variable. To address these challenges, we pioneered a CRISPR/Cas9-based approach that eliminates the need to create a floxed *Kdm8* allele while allowing head-to-head comparisons between *Kdm8*-sufficient and knockout malignant cells within the same mouse. Thus, several unique conclusions can be drawn from the features of our GEM model. First, our data confirm that cellular plasticity is likely shaped by extrinsic factors, such as the hypoxia-Kdm8 axis. Second, hypoxia appears to be a strong predictor of *Kras* imbalance, which can be attributed to the loss of the *Kras^WT^* allele and/or the amplification of the oncogenic *Kras^G12D^* allele. Similar to human PDA ^83^, our FISH analysis supports arm-level *Kras* copy number gain as a primary cause of the allelic imbalance observed in this GEM model. Third, stochastic depletion of *Kdm8* in the malignant cells within a tumor enabled major transcriptomic and histological changes that are associated with *Kdm8* loss. These changes include an almost invariant loss of differentiation and Vimentin expression upon *Kdm8* ablation in malignant cells juxtaposed with Kdm8-expressing neighbor cells, a feature simply not available in conventional *KPC* mice.

The impact of epigenetic dysregulation on genome integrity remains largely unexplored. Our findings provide a crucial piece to the puzzle, demonstrating that hypoxia promotes CIN through its regulation of epigenetic modifiers. Previously, Whetstine and colleagues showed that hypoxia induces transient copy-number gains at specific genomic sites by stabilizing KDM4A ^37^. Mechanistically, hypoxia facilitates the recruitment of KDM4A to H3K4 methylated genomic sites, including *EGFR*, by modulating multiple histone lysine methyltransferases and demethylases ^104^. In this context, hypoxia induces transient copy-number gains at specific loci by increasing the demethylase activity of KDM4A, which promotes re-replication of the amplified regions during the S phase ^105^. In contrast, our model supports the hypoxia-induced suppression of KDM8 demethylase activity, which instead promotes a global epigenetic re-wiring, CIN, and stable polyploidy in select chromosomes. However, the mechanisms through which *Kdm8* deficiency leads to unique mitotic defects, such as multipolar spindle formation, remain poorly understood. One possibility is that dysregulated H3K36me2 near centromeres results in reduced compaction during mitosis, which has been proposed to induce genomic instability ^106^. Alternatively, Kdm8 may have other non-histone substrate(s), such as tubulin, where post-translational modifications like methylation play crucial roles in maintaining mitotic fidelity ^107^. Given the complexity of cellular adaptations to hypoxia, it also remains to be determined whether additional O_2_-sensing epigenetic regulators play crucial roles in promoting CIN.

## Methods

### Autochthonous Tumor Model

All *in vivo* experiments were performed in accordance with the Rutgers University Institutional Animal Care and Use Committee (IACUC) guidelines. The *PTC* mice (*Trp53^flox/flox^*;*R26^LSL-Tomato^*;*H11^LSL-Cas9^*, courtesy of Monte M. Winslow) ^58^ were outbred to the *C57BL/6J* background. To initiate PDA in the *PTC* mice, retrograde pancreatic ductal injection of engineered AAV was performed as reported previously ^57^. Briefly, mice were anesthetized prior to laparotomy and bowel displacement. The duodenum and a portion of the pancreas were gently repositioned to visualize the ampulla of Vater - the injection site where the common bile duct enters the duodenum. A microclip (Micro Vascular Clip RS-6472, Roboz) was applied to the common bile duct near the gallbladder base to increase intraductal pressure. A 30-gauge needle was then inserted at a shallow angle into the bile duct, while a serrated curved forceps stabilized the duodenum. Under maintained pressure, 50-100 µL of engineered AAV was slowly infused into the pancreas over 2-3 minutes. Following injection, the pancreas and intestines were returned to the abdominal cavity. The abdominal muscle was sutured, and the skin was closed using surgical staples. Postoperative analgesia was provided using Ethiqa XR (Fidelis Animal Health). Humane endpoints criteria included signs of pain, distress, or moribund status due to PDA tumor burden. Mice that survived beyond one year post-surgery without evidence of PDA development were excluded from the study.

### Transplanted Tumor Models

PDA cells were washed three times in cold PBS before resuspension in 100% Matrigel (356231, Corning) and kept on ice. For subcutaneous tumor studies, 5×10^5^ cells were injected into the dorsal flank of the *NSG* mice. For orthotopic pancreatic tumor studies, cells were resuspended in 100% Matrigel at a concentration of 4×10^6^ cells/mL prior to the procedure. Briefly, mice were anesthetized before a small incision was made to expose the pancreas tail and the spleen. The resuspended PDA cells were injected into the pancreas tail to form a blister (∼25μL) and allowed to reach body temperature and solidify. The pancreas was then placed back into the abdominal cavity before suturing the abdominal muscle and closing the incised skin with surgical staples. For all *in vivo* studies, mice were sacrificed upon reaching the endpoint tumor volume (∼1cm^3^) or morbidity and dissected for fluorescence microscope imaging and histology. Metastases were quantified using custom FIJI macro scripts. Both male and female *NOD/SCID/γc* (*NSG*, 005557, Jackson Lab) mice were used. In the orthotopic tumor studies using the 703T and 606T cell lines, male *C57BL/6J* (000664, Jackson Lab) mice were used to match the PDA cell background.

### Immunoblot Analysis

Cell lysates were prepared using 2x Laemmli buffer (histone 3) or RIPA buffer supplemented with protease inhibitors (P8340, Millipore Sigma). PAGE separated proteins and subsequently transferred onto an Immobilon PVDF membrane (1620177, Millipore Sigma). Membranes were incubated with primary antibodies (Kdm8: NBP1-77074, Novus Biologicals; β-Actin, 4967S, Cell Signaling; total Histone 3, 4499T, Cell Signaling; H3K27me3, 9733T, Cell Signaling; H3K36me2, ab9049, abcam; HIF-1β/ARNT, 5537T, Cell Signaling; H3K27Ac, ab4729, abcam; FLAG, F3165, Sigma Millipore) in 5% skimmed milk overnight at 4°C, followed by three washes with TBST (1% Tween20), and stained with horseradish-peroxidase-conjugated secondary antibodies. After additional washes, chemiluminescence was performed to visualize proteins of interest. Protein densitometry was performed using FIJI.

### Construction of the AAV-Barcode Library

A previously generated AAV-KrasHR2.1Kb-U6-sg*Kras*-PGK-Cre vector ^58^ and pAAV-CAG-GFP (37825, Addgene) were used to generate the backbone of the AAV library (sg*Kras*: 5’ GACTGAGTATAAACTTGTGG). An additional U6-sgRNA cassette targeting *Kdm8* or non-targeting control (sg*Kdm8*: 5’ tcaggtgctgatcatgtcag; sg*NT*: 5’ gcgaggtattcggctccgcg) was subcloned into the AAV vector from the Lenti-CRISPRv2 vector (52961, Addgene) using the NEBuilder HiFi assembly kit (E2621, NEB). The resulting AAV-KrasHR2.1Kb-hU6-sg*Kdm8*-U6-sg*Kras*-PGK-Cre vector was then validated for ITR integrity before HiFi assembly using the *de novo* AvrII and BsiWI sites to build the barcoded *Kras^G12D^* library as reported ^58^. Two separate oligo pools with a predicted diversity of 2,359,296 synonymous mutations (mostly at wobble bases) on both sides of the oncogenic *Kras^G12D^*mutation were generated from IDT (oPool, IDT) and PCR amplified using the flanking sequences that are homologous to the linearized AAV vector. In addition, we incorporated 4 additional synonymous mutations at *Kras* codon 3, 4, 5, 6, 7, and 8 to confer sg*Kras* resistance (GAGTATAAACTTGTGGTG to GAaTAcAAgCTaGTaGTc). The pooled oligos and the linearized AAV vector were assembled overnight, ethanol precipitated, and electroporated into Stbl4 electrocompetent cells (11635018, Thermo Fisher Scientific) to maintain a representation of 20.84 colonies per *Kras^G12D^* barcode. The assembled AAV vectors were then validated through Illumina sequencing and packaged into the AAV library (AAV8, SignaGen).

### Genomic Cleavage Detection Assay

Targeted indel formation was determined using the GeneArt Genomic Cleavage Detection Kit (A24372, Thermo Fisher Scientific). Briefly, PCR was performed using the genomic DNA isolated with the DNeasy Blood and Tissue Kit (Qiagen) and primers flanking the *Kdm8* target region. The PCR amplicon was purified using the NucleoSpin PCR clean-up kit (Takara) and subjected to slow re-annealing using a thermocycler. The heteroduplex DNA containing indels was subsequently detected using an endonuclease and resolved on a 2% agarose gel.

### Quantitative PCR

Cells were homogenized using TRIzol (Invitrogen) to release the total RNA and quantified by a Nanodrop spectrophotometer before conversion to cDNA with the High-Capacity cDNA Reverse Transcription kit (43-688-14, Thermo Fisher Scientific). Transcript quantification was performed using PowerUp SYBR™ Green Master Mix (Thermo Fisher Scientific).

### Histopathology and IHC

Primary PDA tumors, lungs, and livers were dissected from tumor-bearing mice at the endpoint and fixed in 4% formalin and paraffin embedded. Hematoxylin-eosin staining was performed with a Sakura Prisma Hematoxylin and Eosin stainer according to the manufacturer’s protocol (Sakura Finetek USA). Antibodies for Kdm8 (101664-T40, SinoBiological), Vimentin (5741, Cell Signaling), Tomato (600-401-379, Rockland), and Cytokeratin 19 (TROMA-III, Developmental Studies Hybridoma Bank) were used in IHC on a Roche Discover Ultra Immunostainer (Roche).

### Immunofluorescence and the Analysis of Kdm8 and Galectin-4 in PDA Tumors

Sections were first deparaffinized on the instrument and subjected to buffer washes. Antigen retrieval was performed using a pH 9 retrieval buffer for 76 minutes at 95 °C. An inhibitor block was then applied to reduce nonspecific binding and inhibit endogenous enzymatic activity. The primary JMJD5 antibody (1:700) was manually applied for 60 minutes at 37 °C. After a buffer wash, OmniMap anti-Rabbit secondary antibody (760-4311, Roche) was applied at room temperature, followed by the addition of Rhodamine 6G (760-244, Roche). Antibody denaturation was performed followed by manual application of the primary Galectin-4 antibody (1:4000, PA5-34913, Fisher Scientific) for 60 minutes at 37 °C. The OmniMap Rabbit secondary was re-applied, followed by the addition of Cy5 (760-238, Roche). After denaturation, the anti-RFP antibody (1:3500) was then manually applied for 60 minutes at 37 °C, followed by the addition of OmniMap Rabbit and FAM (760-243, Roche). DAPI (760-4196, Roche) was used for nuclear staining. Four-channel fluorescence images were acquired using an Olympus® VS120 whole-slide fluorescence scanner equipped with a 20x objective. Image analysis was performed using the Visiopharm® digital pathology analysis platform. Regions of interest (ROIs) were manually annotated, and automated cell segmentation was performed based on DAPI nuclear staining. Quantification of fluorescence signal intensity was conducted on a per-cell basis within the annotated ROIs using a custom R code.

### Quantification of Kdm8 and Vimentin in PDA Tumors

Kdm8, Vimentin, and Tomato IHCs were performed on serially sectioned slides and imaged at 20x magnification on an Olympus VS-120(R) whole slide scanner. Each set of whole slide image stack was digitally aligned with the VisioPharm(R) digital pathology analysis suite. The aligned images were divided into thousands of rectangular regions per sample with 354 microns in width and height, and the regions of interest were defined as having more than 40% of the normalized Tomato signal. Normalized expression and the correlation between Kdm8 and Vimentin across all regions of interest were quantified.

### Human PDA Data and Analyses

Bulk RNA-Seq data from the PanCuRx study ^38^, ICGC Controlled Data ^108^, TCGA (unrestricted access through cBioPortal) ^70^, and Dhara et al ^71^ were acquired through granted access to the respective data portals. Processed snRNA-Seq data from Hwang et al was accessed through the Broad Institute Single Cell portal ^73^ (https://singlecell.broadinstitute.org/single_cell). The PanCuRx study was conducted with support of the Ontario Institute for Cancer Research (PanCuRx Translational Research Initiative) through funding provided by the Government of Ontario, the Wallace McCain Centre for Pancreatic Cancer supported by the Princess Margaret Cancer Foundation, the Terry Fox Research Institute, the Canadian Cancer Society Research Institute, and the Pancreatic Cancer Canada Foundation. For GSEA, genes were ranked by DESeq2-derived log_2_ fold change between all basal-like and classical-like samples using raw read counts from the PanCuRx and TCGA data sets and the enrichments of hallmark and custom gene signatures were quantified. Kaplan-Meier survival of the KDM8-induced gene signature score high (>IQR Q2 value) and low PDA patients from the PanCuRx data set was performed using ‘rms’ and ‘survival’ packages in r. To quantify the significance of association between the status of liver metastasis at diagnosis and the following independent variables, including tumor size (scaled not centered), classical and basal subtype gene signatures, hallmark hypoxia, KDM8-induced gene signature, and Eastern Cooperative Oncology Group performance status (ECOG) in the PanCuRx cohort, we performed univariate logistic regression using the ‘pROC’ package. To quantify the association between the *KRAS* allelic imbalance (ordinal variables 0-3) and independent variables including the anatomic location (liver metastasis=1, primary PDA=0), classical and basal subtype gene signatures and state (classical=1, basal=0), hallmark gene signatures (n=50), and KDM8-induced and suppressed gene signatures, we performed univariate ordinal logistic regression to derive the coefficients, confidence intervals, and nominal p-values using the ‘ordinal’ package in r. All gene signatures were scaled using scale in r (center = T, scale = T).FDR-adjusted p-values were derived for the correction of multiple comparisons.

### Malignant Cell Isolation from the *PTC* Mice

Tumors and metastases harvested from the *PTC* mice were dissected, minced, and digested to facilitate tissue dissociation as previously described ^11^. Briefly, dissected tumors were dissociated in a digestion buffer (HBSS-free with collagenase IV, dispase, protease inhibitor cocktail) at 37°C for 30 min followed by quenching with pre-chilled quench solution (L-15 medium, FBS, DNase). Dissociated cells were passed through a 40μm strainer and resuspended in FACS buffer. Malignant cells in the ascites were prepared through ACK lysing buffer incubation at room temperature for 1-2 min prior to straining and FACS buffer resuspension. In most cases, a small aliquot of dissociated cells was dispensed on a 10cm plate for the establishment of parental cell lines. The *ex vivo* isolated cells were subsequently stained with DAPI for viability and antibodies to the lineage markers including CD45 (103111, Biolegend), CD31 (102409, Biolegend), Ter119 (116211, Biolegend), and F4/80 (123115, Biolegend) for the exclusion of most non-malignant cells. Pure, viable, Tomato-positive, DAPI-negative, and lineage-negative malignant cells were isolated using a BD Biosciences Influx High Speed Cell Sorter at the Flow Cytometry Core Facility of Rutgers Cancer Institute.

### Single-Cell RNA-Seq Library Preparation

Using the *ex vivo* sorted cells, Single-cell RNA-seq libraries were prepared using the 10X 5’ v2 dual indexed kit according to the user guide. In 3 metastatic lesions, we spiked in different numbers of Jurkat cells due to low cell counts of sorted Tomato-pos malignant cells (A252, A147, and L254). Following cDNA amplification, the cDNA pool was split and used for scRNA-Seq library construction, Kdm8-Seq, and KrasBC-Seq.

### ChIP-Seq Preparation and Sequencing

Chromatin immunoprecipitation (ChIP) sequencing was performed as reported previously ^109^. Briefly, 2.5×10^7^-1×10^8^ 688M cells were cultured for each ChIP. To crosslink, formaldehyde solution was directly added to cell culture media to a final concentration of 1% for 10 min, followed by quenching with 125mM glycine solution. Fixed cells were rinsed twice in ice-cold PBS and lysed in ice-cold lysis buffer (50mM HEPES-KOH, pH 7.5, 140mM NaCl, 1mM EDTA, 10% Glycerol, 0.5% NP-40, 0.25% Triton X-100, with protease inhibitor cocktail). Lysates were transferred to a 1.5 ml TPX tube (Diagenode) and sonicated using Bioruptor (Diagenode) three times of 10 min (output level: high, interval: 0.5). An aliquot (50 μl) of the lysates was reserved for QC and input library. For immunoprecipitation, 4μg of ChIP-grade antibodies against H3K36me2 (ab9049, abcam), H3K27me3 (ab192985, abcam), or H3K27ac (ab4729, abcam) was added to the lysate (containing ∼50μg of sheared gDNA) and rotated overnight at 4°C, followed by an incubation with 25 μl of Protein A Dynabeads (Life Technology) for 2 hours at 4°C. The beads were washed three times with 800 μl ice-cold LiCl washing buffer (100 mM Tris-HCl, pH7.5, 500 mM LiCl, 1% NP-40 and 1% sodim deoxycholate) prior to elution and reverse crosslink through an incubation with 400 μl of digestion buffer (0.1 M Tris-HCl, pH 7.5, 100 mM NaCl, 50 mM EDTA, 1% SDS, and 200 μg/ml of proteinase K) at 65°C for 2 hours with agitation every 30 min. The Input lysate was treated the same way to reverse crosslink. Eluted DNA samples were purified by organic extraction and ethanol precipitation. DNA libraries were prepared using KAPA Hyperprep Kit (Roche) and sequenced on the Illumina Nova-seq X platform. For ChIP-qPCR, primers were designed such that (i) amplicon size was <100bp amd (ii) within regions that demonstrated significantly elevated signals following *Kdm8* knockdown. Input DNA was diluted 1:200 and ChIP DNA was diluted 1:20 for the qPCR reaction performed using PowerUp SYBR™ Green Master Mix (Thermo Fisher Scientific).

### ATAC-Seq Library Preparation and Sequencing

Libraries were generated using an updated method ^110^. Briefly, viable 688M PDA cells were first treated with DNase in HBSS for 30 min at 37°C prior to nuclei release in pre-chilled ATAC resuspension buffer (Tris-HCL pH 7.4, digitonin, Tween-20, NP40). Transposition was carried out in transposition mixture (1xTD buffer, 100nM Tn5 transposase, digitonin, Tween-20) for 30 min at 37°C in a thermomixer with 1000 rpm. Transposed fragments were cleaned up with a Zymo DNA Clean and Concentrator-5 Kit. Partially amplified libraries were first quantified by qPCR to determine additional cycles needed and fully amplified using the NEBNext 2x MasterMix. Libraries were sequenced on an Illumina NextSeq 2000 sequencer.

### Library Preparation for Targeted Sequencing

The preparation for targeted KrasBC-Seq and Kdm8-Seq was based on a modified 10X VDJ immune profiling protocol. Briefly, amplified 10X cDNA libraries were used as templates for nested PCR reactions with none adapted primers for PCR I (12 cycles for KrasBC-Seq and 13 cycles for Kdm8-Seq) and composite primers with sample indices (denoted as “N”) and Illumina adapter sequences for PCR II (10 cycles for KrasBC-Seq and 11 cycles for Kdm8-Seq; KrasBC-Seq primers, forward: 5’ aatgatacggcgaccaccgagatctacactctttccctacacgacgctc; reverse: 5’ caagcagaagacggcatacgagatNNNNNNNNgtgactggagttcagacgtgtgctcttccgatctgtaggagtcctctatcgtagggtc; Kdm8-Seq, forward primer is the same as KrasBC-Seq, reverse: 5’ caagcagaagacggcatacgagatNNNNNNNNgtgactggagttcagacgtgtgctcttccgatctagctcctcctctttgtcaggc). Both PCR reactions were performed using ExTaq polymerase (Takara) and the amplicons were purified and size-selected using SPRIselect Beads (Beckman Coulter). Beads purified PCR amplicons from PCR II were quantified on a Qubit HS DNA quantification kit and a BioAnalyzer (Agilent) prior to sequencing (Novogene).

### Target Gene Knockdown, Knockout, and Re-expression in PDA Cell Lines

To knockdown a gene in PDA cells, shRNA sequences were designed and subcloned into the TRC pLKO.1 vector (10878, Addgene; sh*GFP(Control)*: 5’ gcaagctgaccctgaagttcat; sh*Scramble(Control)*: 5’ cctaaggttaagtcgccctcg; sh*Kdm8-1*: 5’ cgcacattcttcataacacca; sh*Kdm8-2*: 5’ gctcctgatgtcatgttagag; sh*Kdm2a*: 5’ gctccaaaccaacaaatataa; sh*KDM8-1*: 5’ cctgttcatcccggtgaaata; sh*KDM8-2*: 5’ tcagcaaatacatcgtgaatg; sh*KDM8-3*: 5’ cgaggtacacagatgaggaat; sh*Nsd2*: 5’ ccagaaagagcttggatattt; sh*Smad4*: 5’ gccagctacttaccatcataa). To knockout *Kdm8* or *Arnt* (sg*Kdm8-1*: 5’ caaggtacacagatgaagac, sg*Kdm8-2*: 5’ atgtactgcagactacgaga; sg*Kdm8-3*: 5’ tcgtagtctgcagtacatcc; sg*Arnt-1*: 5’ gacatcagatgtaccatcgc; sg*Arnt-2*: 5’ agggtttccagaagcaatgg), we subcloned sgRNA sequences into the PX458 vector and transiently transfect PDA cells using TransIT-293 transfection reagent (MIR2704, Mirus Bio). Single GFP+ cells were sorted onto 96-well plates 48 hours post transfection to establish knockout clones and validated using western blot. To re-express *Kdm8* cDNA, murine Kdm8 coding sequence was subcloned into the Lv241 lentiviral vector using TOPO TA cloning kit (450641, Thermo Fisher Scientific). *Kdm8* variants were subsequently generated using Q5 site-directed mutagenesis kit (E0554, NEB). *Kdm8* knockout PDA clones were infected with lentiviral vectors carrying *Kdm8* variants and selected with puromycin or blasticidin.

### Lentivirus Production and Infection

Lentivirus were generated by co-transfecting the viral shuttle vector with Delta8.2 and VSV-G packaging plasmids into the 293T cells using polyethylenimine (23966-1, Polysciences). Medium was replenished 24 hours post transfection and supernatants were collected at 48 hour, frozen, thawed, and spun down at 2000 rpm before infecting PDA cells.

### Bulk RNA-Seq Library Preparation and Analysis

Total RNA was isolated from PDA cells using RNeasy Mini kit (Qiagen) prior to library construction using the NEBNext Ultra II RNA Library Prep Kit for Illumina (NEB). Libraries were sequenced on an Illumina NovaSeq 6000 sequencer (paired-end 2×150bp, Novogene). Reads were mapped to the mm39 genome build using the STAR aligner (v2.7.5a). Reads counts at both transcript and gene levels were derived using htseq-count (--stranded=no --type=exon --mode=intersection-nonempty) and transcripts per kilobase of exon and scaled in million or TPM was derived using custom r scripts and the mm39 reference gtf file. Differentially expressed genes, fold change, and adjusted p-values were derived from read counts using DESeq2 (v1.46.0) ^111^. Genes with low abundance were removed from further analysis. For GSEA (Broad Institute), log_2_ fold change ranked gene lists were assessed for enrichment of hallmark (v7.5.1), curated gene sets (c2.all.v7.5.1.symbols, n=4,133), or custom gene signatures.

### *In Vitro* Cell Count and Viability Assays

Cells were seeded in triplicate into a 6-well plate at a density of 5×10^4^ cells/well on day 0. The plates were then cultured under normoxia or hypoxia for 96 hours. On day 2 and day 4, the cells were trypsinized and total cell count was determined using a Vi-CELL XR Cell Viability Analyzer (Beckman Coulter). For viability, cells were seeded onto a 96-well plate at a density of 2.5-10×10^3^/100μL/well and allowed to attach overnight. Cells were then treated with the Kras inhibitor for 48 hours before equal volumes of CellTiter-Glo (Promega) were added and mixed on an orbital shaker for 3 minutes. Luminescence was quantified using a plate reader (Tecan).

### Single-cell Transcriptomic Pipeline

Fastq files were mapped to a custom mm39 genome build (GRCm39 release 108 with the Tomato transcript) using the 10X CellRanger pipeline (v6.0.2). To exclude Jurkat cells in samples A252, A147, and L254, reads were also mapped to the GRCh38 genome reference. We used Seurat (v4) and custom R scripts for the identification of cell clusters, differential genes, and all the downstream cell state analyses ^112^. Briefly, CellRanger output sparse matrices were merged and read counts were normalized after removing potential doublets, dying cells, and Jurkat cells based on GRCm39 transcript count, mitochondria percent, and GRCh38 transcriptomic features, respectively. Distinct clusters of malignant cells were uncovered using SNN modularity optimization-based approach following linear transformation of data and dimensionality reduction. To calculate gene signature scores, we performed single sample GSEA using the GSVA r package (v4.4.2). For RNA velocity analysis, we inferred the latent time and estimated differential kinetics using scVelo ^81^ (v0.3.3). Briefly, mapped bam files from the CellRanger pipeline were used to generate spliced and non-spliced expression matrices using default velocyto run10x command. Preprocessing was performed to compute first- and second-order moments for velocity estimation and subsequently the full splicing kinetics using the scv.tl.velocity() function. We then projected the velocity or latent time estimates onto the UMAPs for visualization.

### *Kras^G12D^* Barcode Calling for Clonal Identity

To quantify the expression of wildtype and barcoded *Kras^G12D^* transcripts in each cell, KrasBC-Seq 3’ reads that passed QC were first trimmed using cutadapt (v3.4, -g “ctctatcgtagggtcata;max_errors=0.2;optional…cattttcagcaggcct;max_errors=0.2”). Trimmed reads were subsequently mapped to a custom reference containing wildtype *Kras* and the engineered *Kras^G12D^* BC sequences using bowtie2 (v2.4.1). Additionally, the paired reads were subjected to the CellRanger pipeline to resolve UMI and the 10X barcode (cellBC) identities. Mapped reads were collapsed with an identical cellBC, *Kras^G12D^* BC (or wildtype *Kras*), and UMI combination into a unique transcript in each cell. Owing to the extremely low incidence of homology-directed repair that is anticipated to occur twice in a cell ^58^, a vast majority of malignant cells should only carry one *Kras^G12D^* barcode. Based on this assumption, we quantified the UMI counts of all *Kras^G12D^*BCs with edit distance less than 3 and assigned the dominant *Kras^G12D^*BC to each unique cellBC. Non-malignant (somatic) cells were defined as having more than one copy of wildtype *Kras* and no detectable *Kras^G12D^*BC. Cells without a dominant *Kras^G12D^* BC and wildtype *Kras* were excluded from downstream lineage analyses. The abundance of *Kras* in a cell quantified by KrasBC-Seq was defined as the sum of unique UMI counts of *Kras^G12D^* BC and wildtype *Kras* normalized against the count of the entire transcriptome and scaled according to sequencing depths. To create the Sankey diagrams, networkD3 (v0.4) was used in custom r codes.

### Kdm8-Seq to Infer *Kdm8* Genome Editing

Similar to the KrasBC-Seq pipeline, Kdm8-Seq 3’ reads were first trimmed using cutadapt to remove the first 26 bp. Trimmed 3’ reads were subsequently mapped to the *Kdm8* transcript (NM_029842) using the bwa aligner (v0.7.17-r1188). To include all malignant cells from the primary PDA tumors, cells without detectable Kdm8-Seq signal were imputed as having zero read count. UMI and cellBC identities in the paired 5’ reads were uncovered using the CellRanger pipeline. Reads were then collapsed into unique UMI and cellBC combinations. The abundance of *Kdm8* in a cell or cellBC quantified by Kdm8-Seq was defined as the unique UMI count normalized against the count of the entire transcriptome and scaled according to sequencing depths.

### ChIP-Seq and ATAC-Seq Data Analysis

Both ChIP-Seq and ATAC-Seq reads that passed the QC process were aligned to a mm39 genome build using bowtie2 (v2.4.1). Bam files were filtered for reads mapped to blacklisted regions and duplicated reads. For the ATAC-Seq pipeline, insertion bed files were made with bedtools (v2.25.0) accounting for the +4/−5 bp offsets by Tn5 ^113^. Merged peaks were called using MACS3 (v3.0.0a6) to derive broad and narrow peak files by comparing ChIP pulldown signals with input samples. Only reads with low false discovery rate (qValue > 5) were included in the analysis. Counts of the pulldown or Tn5 insertions within merged peaks were derived using bedtools and data from technical replicates were used to derive differential change and adjusted p-values using DESeq2, normalized against size factors that reflected mapped read depths. Peaks were annotated using Homer annotatePeaks.pl (v5.1) with the mm39 reference. To quantify the enrichments of transcription factor binding motifs within differential regions, we performed Homer findMotifsGenome.pl (-size given) using curated collection of mouse motifs from the CisBP database (n=7,000 motifs) and comparable numbers of constitutive regions as background. To compare ChIP profiles across samples, mapped reads from replicates were first merged with samtools (v1.3.1) and normalized against comparable input samples using bamCompare (v3.5.4.post1). Aggregate ChIP profiles within differentially regulated H3K36me regions were created using deeptools (v3.5.4.post1).

### Bulk and Single-cell Whole Genome Sequencing and Analysis

For bulk WGS, genomic DNA were extracted using the Qiagen DNeasy Blood and Tissue Kit (69504, Qiagen). The libraries were prepared using the Rapid Plus DNA Lib Prep Kit (ABclonal) prior to sequencing on a NovaSeq X Plus sequencer (Novogene). For single-cell WGS, 688M cells expressing control or a *Kdm8*-targeting shRNA were sorted into a 96-well plate prior to library preparation using the PicoPLEX® Gold Single Cell DNA-seq Kit according to the manufacturer’s protocol (R300670, Takara). To assess the copy number variations, we adopted a previously reported pipeline with some modifications ^114^. Briefly, fastq files from both the bulk and single-cell WGS studies were mapped to an mm39 reference prior to alignments using bowtie2 (v2.4.1). To compute bin boundaries, mm39 fasta files for individual chromosomes were downloaded from UCSC (hgdownload.cse.ucsc.edu/goldenPath/mm39/chromosomes) and used to generate reads from the reference genome. The reads were mapped back to the mm39 reference using bowtie2 and the result was used to generate genomic positions that defined 50,000 bins with equal numbers of mappable positions. Counts of reads within the bins were quantified for each sample with the GC content accounted for. Genomic distribution of WGS signals and the copy number estimation were derived and plotted using custom r codes.

### Cell synchronization and Immunofluorescence microscopy

Cells were seeded in 6-well plates with coverslips and treated with cell culture media containing thymidine (2 mM) for 16 hours, followed by fresh media for 9 hours, then thymidine (2 mM) for another 16 hours. Cells were subsequently released into fresh media for 8-12 hours at anaphase (688 cell lines: 8h; MCF10A cell lines: 12h; 606 cell lines: 8h; Capan-1 cell lines: 10h). Synchronized cells were fixed with 4% paraformaldehyde for 15 minutes and permeabilized with 5% Triton X-100 for 15 minutes. Blocking was performed with 5% goat serum/PBS at room temperature for 1 hour, followed by overnight incubation at 4 °C with primary antibodies diluted in 0.5% goat serum/PBS targeting α-tubulin (Sigma, T9026, 1:1000), γ-tubulin (Sigma, 1:1000), and Pericentrin (Abcam, ab4448, 1:400). Cells were incubated in fluorophore-conjugated secondary antibodies (1:1000 in PBS, anti-rabbit IgG-Alexa Fluor® 488, 4412S; anti-rabbit IgG-Alexa Fluor® 647, 4414; anti-mouse IgG-Alexa Fluor® 488, 4408, all from Cell Signaling) at room temperature for 1 hour and counterstained with DAPI (Fisher, P36935). Images were captured using ZEISS Apotome3 and Nikon A1Rsi Confocal Microscope.

### Metaphase chromosome preparation and Micronuclei quantification

Cells in flasks were made 60%-80% confluent and split a day before harvesting for metaphase chromosomes preparation. Cells were then treated with colcemid (0.1 µg/mL, Invitrogen, 15210-016) at 37°C for 30 minutes. Harvested cells were incubated in a hypotonic solution (0.075 M KCl) at 37°C for 25 minutes, followed by fixation (methyl alcohol/ glacial acetic acid, 3:1 in volume). For micronuclei analysis, fixed cells were dropped onto slides and stained with DAPI. Images of micronuclei were captured and analyzed using a BioView Duet System (Bioview) on an Olympus BX63 fluorescence microscope (Olympus) equipped with a ×60 oil immersion objective (NA 1.3).

### Chromosome painting and FISH analysis

Chromosome painting was performed on fixed cells from metaphase chromosome preparation. Fixed cells were dropped onto slides and baked at 95°C for 5 minutes. Slides were rehydrated in 2 x saline-sodium citrate (SSC, 175.3g NaCl and 88.2g of sodium citrate in 800 mL of ddH_2_O) at 72°C for 2 minutes and then dehydrated in an ethanol series: 70% ethanol, 85% ethanol and 100% ethanol for 2 min each. The custom locus-specific mouse FISH probes for *Kras* were prepared using BAC clone RP23-300B22 (chr6:145163176-145374812, spanning 0.21 Mb), obtained from the BACPAC Resources Center (Children’s Hospital Oakland Research Institute). The BAC probe for *Kras* was generated by nick translation with DY-590-aadUTP (Dyomics) following a previously described protocol ^115^. Similarly, 3 additional custom locus-specific mouse FISH probes targeting chromosome 6 centromere (Cep 6 using BAC clone RP23-341C22 and DY-495-dUTP), chromosome 6 distal (distal chr6 using BAC clone RP24-77K22 and DY-431-dUTP), and chromosome 18 centromere (Cep 18 using RP23-337K12 and DY-530-dUTP) were generated by nick translation and used as reference probes in this study. The 4 color custom FISH probes were mixed at equal ratios and hybridized to metaphases as previously described. Briefly, probes and slides were codenatured at 75°C for 3 minutes on a slide warmer (ThermoBrite System, Molecular Abbott) and hybridized overnight at 37°C in a humidified chamber. Post-hybridization washes were performed in prewarmed 0.4xSSC at 74°C, followed by 2xSSC plus 0.1% Tween at room temperature, serial ethanol dehydration, air drying, and mounting with DAPI antifade VECTASHIELD Antifade Mounting Medium with DAPI (Vector Laboratories) for counterstaining. Images were captured and analyzed by a BioView Duet System (Bioview).

## Acknowledgments

S.G.G. is supported by the National Institute of General Medical Sciences grant R35GM152219. C.M. is supported in part by the National Institute on Aging grant AG068908-02. S.-H.C. is supported by the National Cancer Institute grant 1R01CA285774-01 and in part by the National Institutes of Health grant 1R21CA286389-01. Services, results, and/or products in support of the research project were generated by the Rutgers Cancer Institute Biomedical Informatics Shared Resource, supported, in part, with funding from NCI-CCSG P30CA072720-6852.

## Author contributions

Conceptualization: P.M.G., C.M., S.-H.C. Methodology: P.M.G., K.W.W., Z.Z., W.C., S.G.G., S.-H.C. Investigation: P.M.G., Q.W., Y.-C.C., P.G., R.C., A.K., G.L., S.G.R., Y.M., A.R., K.W.W., L.F., S.B., E.Z., K.P., J.Z.N, S.-H.C. Visualization: Q.W., Y.-C.C., P.G., R.C., A.K., C.M., S.-H.C. Funding acquisition: S.G.G, C.M., S.-H.C. Supervision: S.-H.C. Writing – original draft: S.-H.C. Writing – review & editing: C.M., S.-H.C.

## Materials & Correspondence

All requests for resources, including cell lines, mouse models, plasmids, etc., should be directed to Shin-Heng Chiou through institutional or third-party Material Transfer Agreements (MTA).

## Supplementary information

**Figure S1, related to Figure 1.**
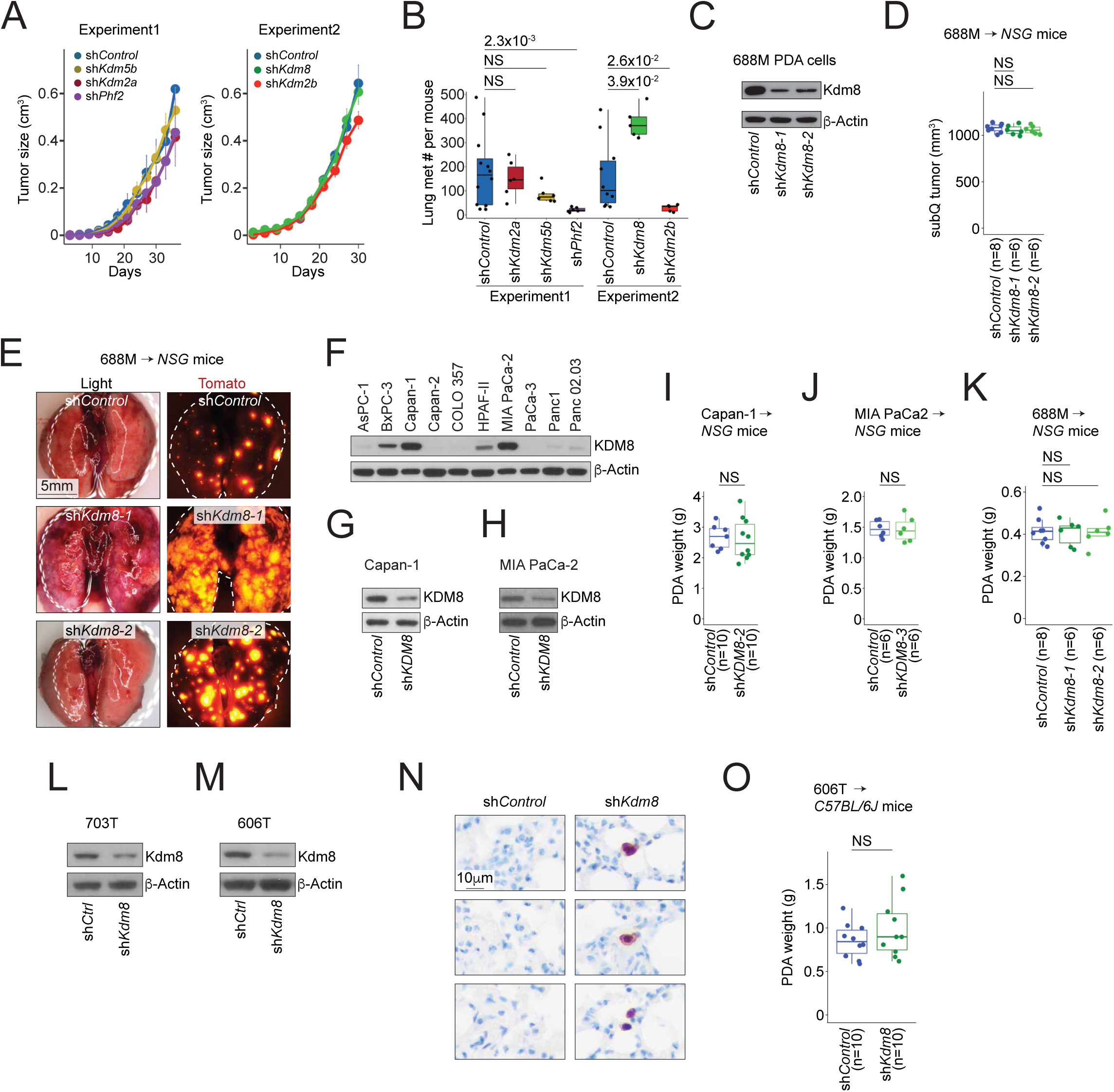
(A,B) Subcutaneous tumor growth (A) and counts of lung metastases (B) seeded from the subcutaneous tumors in the *NSG* mice receiving the Tomato-positive 688M PDA cells transduced with control (sh*Control*, experiment 1, n=12; experiment 2, n=10) or shRNA targeting the indicated *Kdm* genes (sh*Kdm2a*, n=6; sh*Kdm5b*, n=6; sh*Phf2*, n=6; sh*Kdm8*, n=5; sh*Kdm2b*, n=5). Two independent experiments are shown. (C) *Kdm8* knockdown in 688M cells using two independent shRNAs. (D,E) Subcutaneous tumor size (D) and representative light (left) and fluorescent (right) images of the lungs (E) upon sacrifice for the tumor studies in Figure S1B. (F) Immunoblot for the endogenous abundance of KDM8 and β-Actin in the indicated human PDA cell lines. (G,H) *KDM8* knockdown in Capan-1 (G) and MIA PaCa-2 (H) human PDA cells using a *KDM8*-targeting shRNA. (I-K) The primary PDA tumor weight for the orthotopic tumor studies using Capan-1, MIA PaCa-2, and 688M PDA cells. (L,M) *Kdm8* knockdown in 703T (L) and 606T (M) murine PDA cells using a *Kdm8*-targeting shRNA. (N) Representative Tomato IHC images demonstrating individual micrometastatic PDA cells in the lungs of the *B6* mice receiving sh*Kdm8*-expressing 606T PDA cells (right) compared to control (left). (O) The primary PDA tumor weight in each *B6* mouse orthotopically transplanted with 606T PDA cells as in Figures 1N-1P. NS, not significant. Each dot is a mouse and boxes represent medians with interquartile range between 25th and 75th percentiles.

**Figure S2, related to Figure 2.**
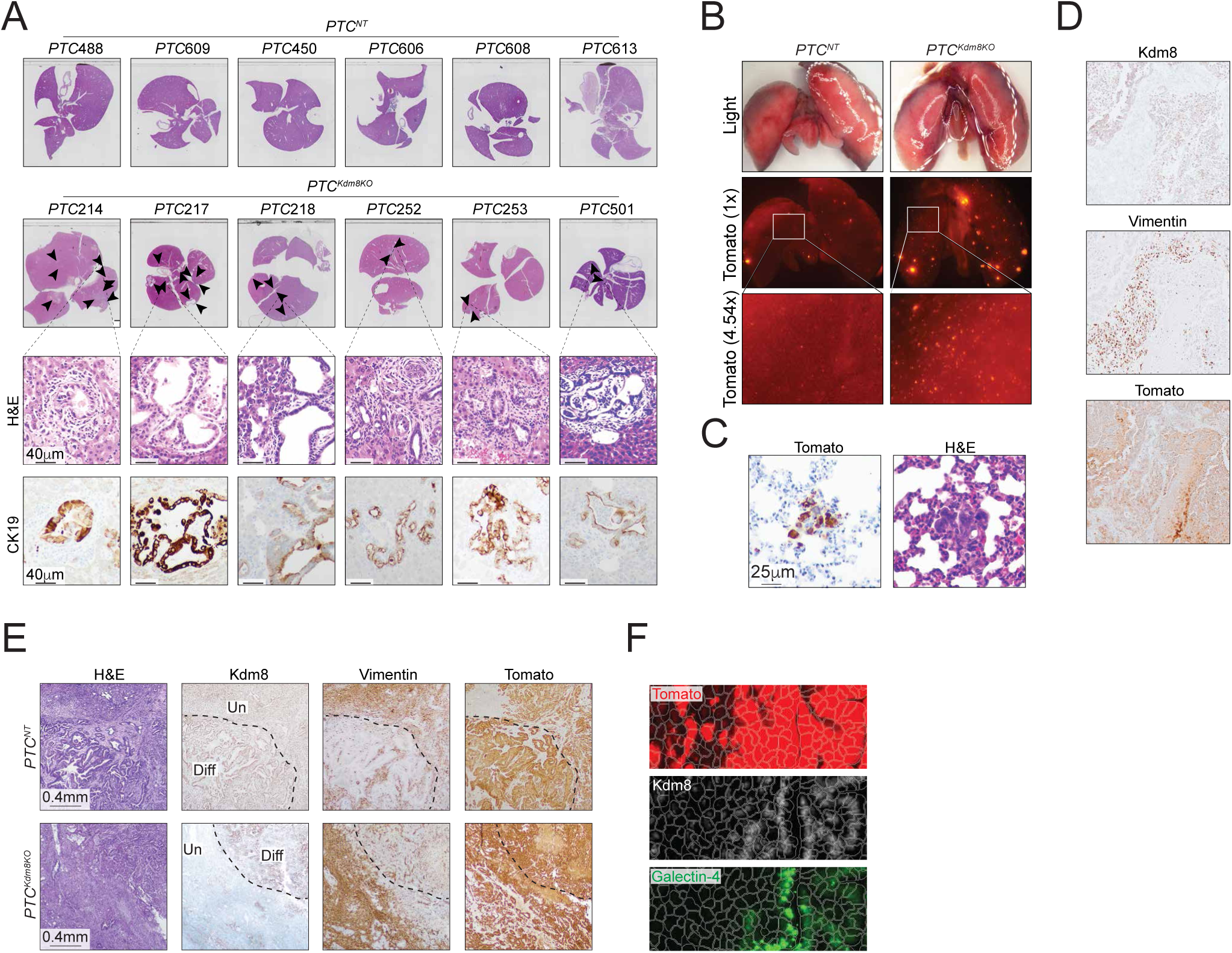
(A) Haematoxylin and eosin staining (H&E) and CK19 immunohistochemistry (IHC) of liver samples from *PTC^NT^* (top row) and *PTC^Kdm8KO^*(2nd, 3rd, and 4th rows) mice. (B) Representative light (top) and fluorescent images of the whole lungs (2nd row) and zoom-in regions (3rd row) from a *PTC^NT^* (left) and a *PTC^Kdm8KO^* (right) mouse. (C) Representative H&E (right) and Tomato IHC (left) of lung micrometastases in a *PTC^Kdm8KO^* mouse. (D) Representative IHC of Kdm8, Vimentin, and Tomato of serial sections from a primary PDA region as in Figure 2M. (E) Representative H&E and IHC of Kdm8, Vimentin, and Tomato of serial sections from primary PDA tumors of a *PTC^NT^* (top) and a *PTC^Kdm8KO^*(bottom) mouse. Dotted lines demarcate the fully differentiated (diff) and poorly differentiated (Un) regions. (F) Representative images as in Figure 2I of a *PTC^Kdm8KO^* tumor with automated cell segmentation, overlaid with pseudocolors for Tomato, Kdm8, and Galectin-4 (top to bottom).

**Figure S3, related to Figure 3.**
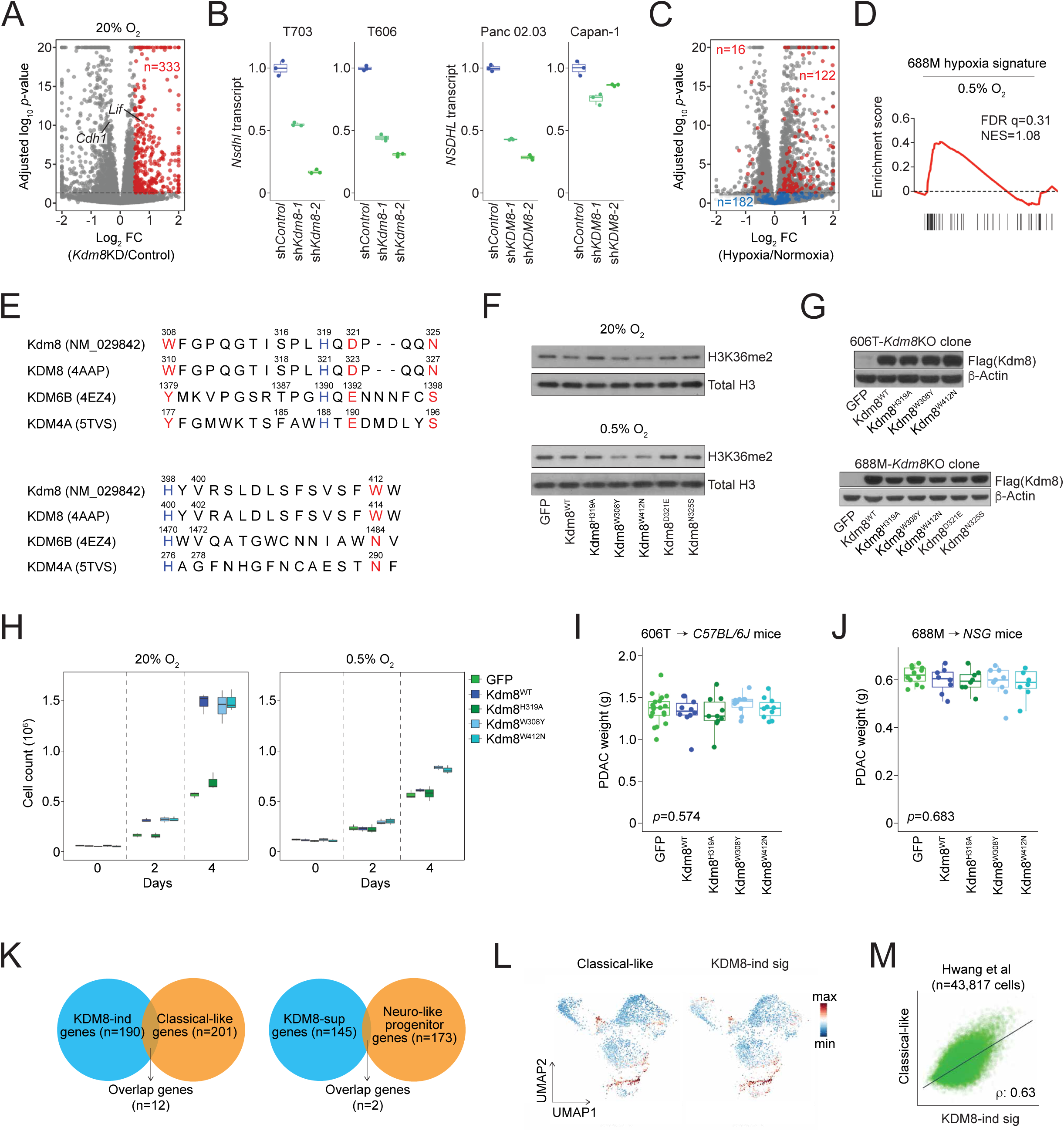
(A) Volcano plot of differential genes induced upon *Kdm8* knockdown in normoxia (n=333 genes upregulated, red). *Lif* and *Cdh1* are highlighted for Figure 5F. (B) Expression of *Nsdhl* and *NSDHL* following *Kdm8* and *KDM8* knockdown the indicated PDA cell lines, as measured by quantitative PCR. (C) Volcano plot of differential genes induced or suppressed by hypoxia in 688M cells. Genes that are induced by *Kdm8* knockdown are highlighted (red, adjusted p-value < 0.05; blue, non-significant under hypoxia). (D) GSEA of the 688M hypoxia gene signature enriched in *Kdm8* knockdown 688M cells cultured in hypoxia. (E) Alignment of the JmjC domains of human KDM8, KDM6B, and KDM4A (with PDB identifiers). Red, non-conserved residues that share consensus in KDM6B and KDM4A; blue, conserved residues among all three KDMs. Murine Kdm8 is shown for comparison. KDM8 residue 328-399 are omitted due to the absence of conserved residues in KDM6B and KDM4A, and their high divergence within KDM8. (F) Immunoblots of H3K36me2 and total H3K36 in *Kdm8* knockout 688M cells re-expressing the indicated Kdm8 variants cultured in normoxia (top) or hypoxia (bottom). (G) Immunoblots of indicated Flag tagged Kdm8 variants in *Kdm8*-knockout 606T (top) and 688M (bottom) clones. β-actin shows equal loading. (H) Cell counts of *Kdm8* knockout 688M cells re-expressing the indicated Kdm8 variants cultured in normoxia (left) or hypoxia (right) following 0, 2, and 4 days post seeding. Comparable results were observed using 606T cells (not shown). (I,J) Pancreatic tumor weights at the time of sacrifice for the tumor studies shown in Figures 3E and 3F, respectively. (K) Venn diagrams demonstrating the shared genes between the KDM8-indued and the classical gene signatures and those between the KDM8-suppressed and the neuro-like progenitor (Hwang et al, 2022 Nat Genet) gene signatures. (L,M) UMAP demonstrating human PDA malignant cells defined in Hwang et al, 2022 Nat Genet colored by the KDM8-induced and classical gene signatures (L). The relationship between the scores of the two gene signatures is shown with the Pearson correlation coefficient ρ (M).

**Figure S4, related to Figure 4.**
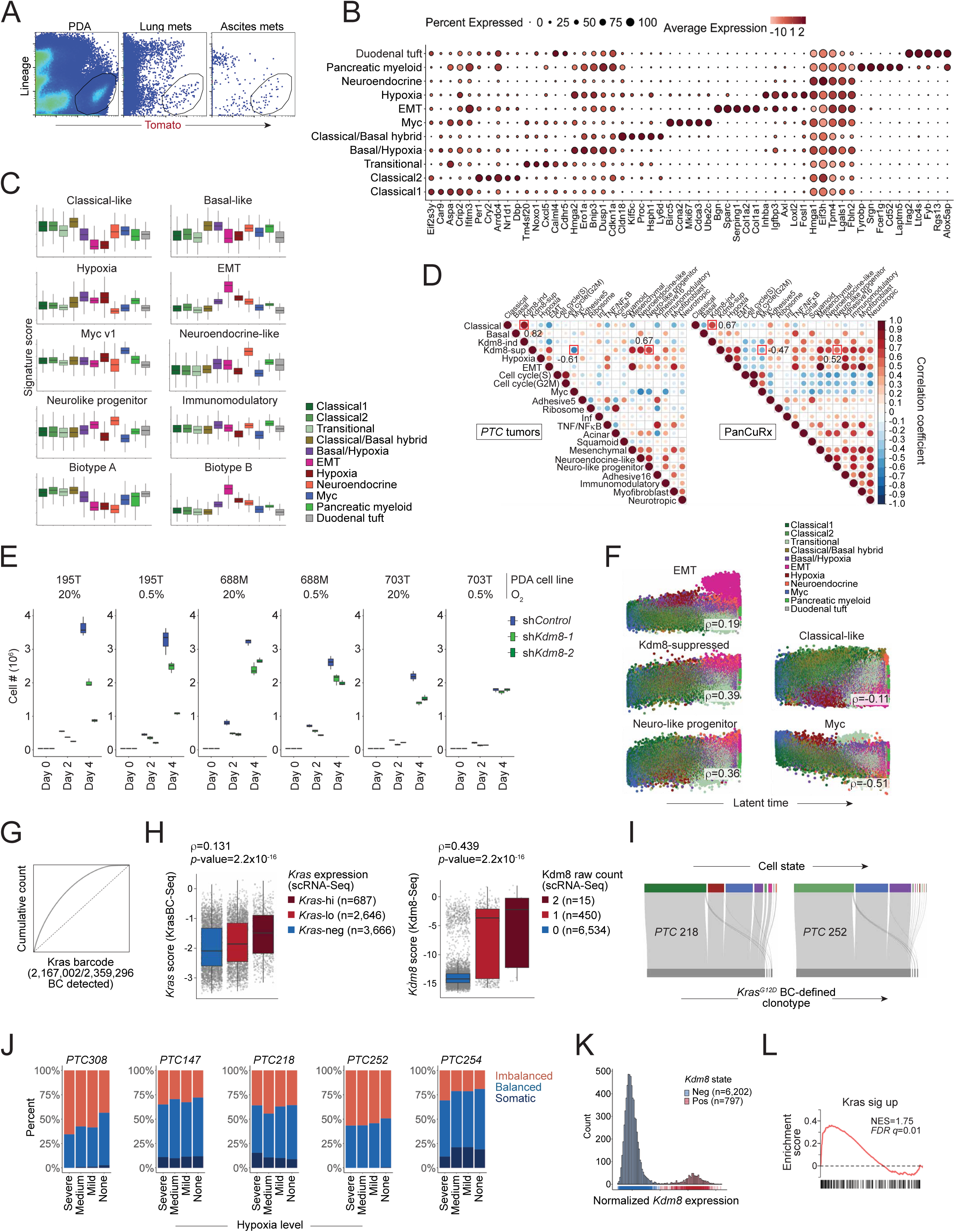
(A) Representative FACS density plots of the Tomato-positive, lineage (Cd31, Cd45, Ter119, and F4/80)-negative malignant cells from the primary PDA tumor, lung metastases, and ascites of a *PTC^Kdm8KO^* mouse. The gate used to sort the Tomato+lineage-cells is shown. (B) Dot plot showing select differential genes across the identified Seurat cell clusters defined in Figure 4B. Dot size represents the percentage of cells expressing the differential genes and color represents the abundance of the differential genes. (C) Relative abundance of the previously reported malignant programs (Hwang et al, 2022 Nat Genet; Di Chiaro, et al, 2024 Cancer Cell; Moffitt et al, 2015 Nat Genet) and GSEA hallmark gene signatures across the 11 clusters of malignant cells defined in Figure 4B. (D) Dot plots demonstrating the Pearson correlation coefficient between any two malignant programs as in C quantified in the current study (left) and the bulk tumors from the PanCuRx PDA cohort (right). Select comparisons are highlighted with their Pearson correlation coefficients annotated. (E) Total cell counts on the indicated days following normoxic or hypoxic culture of 3 murine PDA cell lines expressing control or 2 *Kdm8*-targeting shRNAs. (F) Abundance of the indicated malignant programs plotted along the latent time and colored by the clusters defined in Figure 4B. Pearson correlation coefficients (ρ) are shown. (G) Cumulative plot for the AAV plasmid library. (H) Relationships between the transcript abundance measure by 10x scRNA-Seq (x-axis) and KrasBC-Seq (left, y-axis) or Kdm8-Seq (right, y-axis). Pearson correlation coefficients (ρ) and p-values are shown. (I) Sankey diagrams for all barcoded (bottom) malignant cells isolated from two *PTC^Kdm8KO^* mice that correspond to the malignant programs defined in Figure 4B (top). (J) Proportions of malignant cells isolated from 5 *PTC^Kdm8KO^* tumors, categorized by the indicated *Kras^G12D^*/*Kras^WT^* allelic conformation, across varying levels of hypoxia (x-axis, from none to severe), quantified using the GSVA method. (K) Histogram of *Kdm8* expression in the malignant cells from 5 *PTC^Kdm8KO^* tumors (n=6,999) estimated by Kdm8-Seq. (L) GSEA for the indicated hallmark gene signature (Kras signaling up) enriched in Kdm8^neg^ malignant cells as in Figure 4L.

**Figure S5, related to Figure 5.**
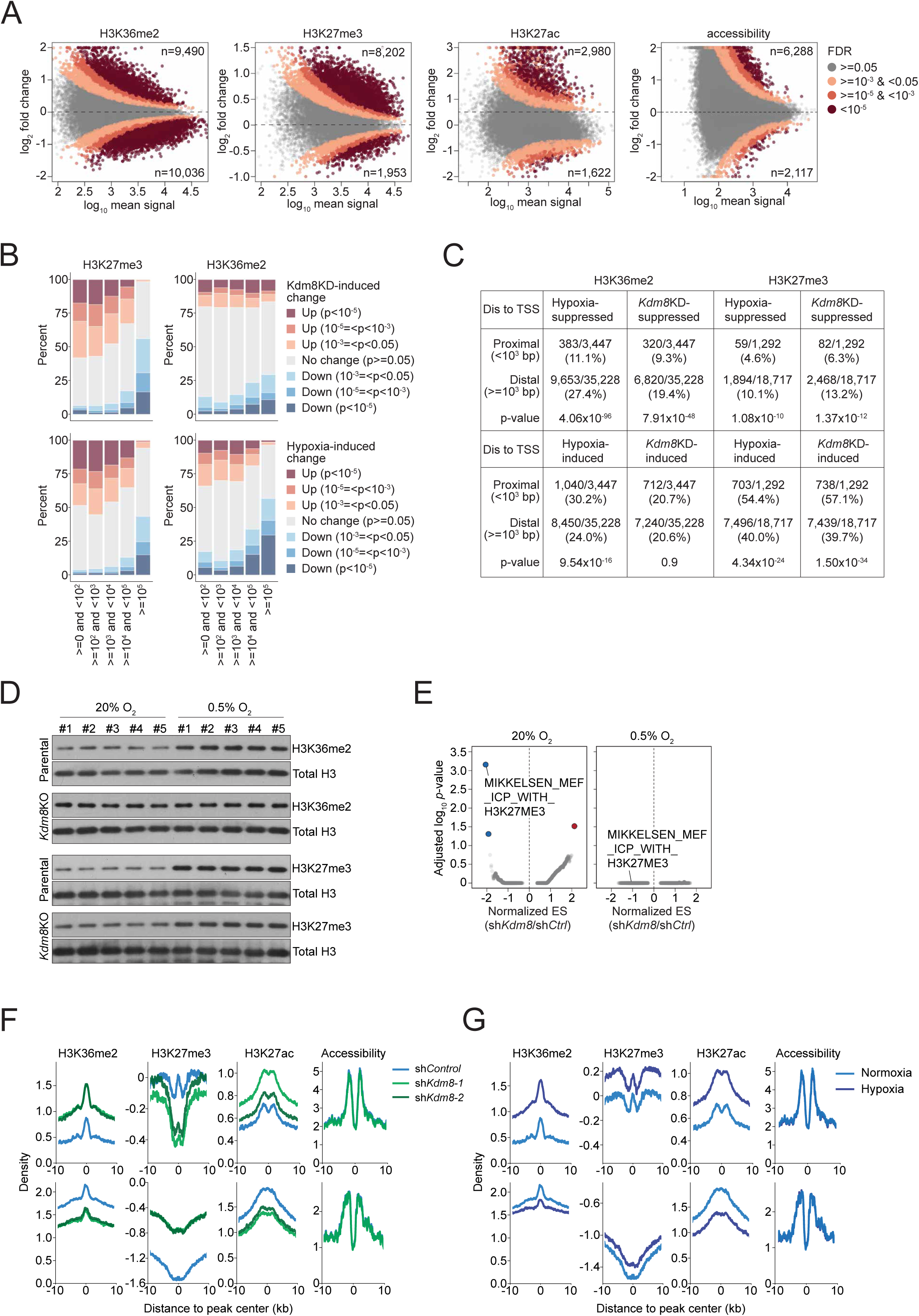
(A) Hypoxia-induced differential H3K36me2, H3K27me3, and H3K27ac ChIP-Seq signals, as well as chromatin accessibility measured by ATAC-Seq (left to right), plotted against the mean read counts per region, as in Figure 5A. Dotted lines represent zero log_2_ fold change (FC), with positive values representing hypoxia-induced increases. FDR (false discovery rate) reflects the probability that a differential region exhibits a significant change induced by hypoxia. The number of significantly altered regions (FDR < 0.05) is indicated at the top and bottom of each plot. (B) Annotations of all H3K36me2 (right) and H3K27me3 (left) regions by Homer’s annotatePeaks.pl, based on their distance to the nearest transcription start site (TSS), grouped into 5 bins with indicated genomic ranges in base pairs (x-axis). The y-axis shows the proportion of regions within each genomic range, stratified by the p-value of *Kdm8* knockdown (*Kdm8*KD)-induced (top) and hypoxia-induced (bottom) changes. (C) Fractions of proximal (<10^3^ bp to TSS) and distal (>=10^3^ bp to TSS) H3K36me2 (left) and H3K27me3 (right) genomic regions that are induced (bottom) or suppressed (top) by hypoxia or *Kdm8*KD. Chi-squared test p-values for the association between distance to TSS and *Kdm8*KD- or hypoxia-induced changes are shown. (D) Immunoblots showing H3K36me2, H3K27me3, and total H3 levels in parental 688M and a *Kdm8* knockout (*Kdm8*KO) clone cultured in normoxia or hypoxia (0.5% O_2_). Data from 5 technical replicates per group are shown. (E) Gene set enrichment analysis (GSEA) of the curated gene sets (C2, v7.5.1) depleted in *Kdm8* knockdown 688M cells cultured in normoxia (left) or hypoxia (right). Normalized enrichment score (ES) for the gene signatures and the corresponding FDR-adjusted p-values are shown. sh*Ctrl*, control shRNA. The MEF cell derived gene signature bearing the H3K27me3 mark (Mikkelsen *et al*, 2008 Nature) is highlighted. (F,G) Aggregate signals of H3K36me2, H3K27me3, and H3K27ac ChIP-Seq, as well as chromatin accessibility measured by ATAC-Seq, within 3,973 genomic regions that are co-induced by *Kdm8*KD and hypoxia for H3K36me2 (top), and 3,838 regions that are co-suppressed by *Kdm8*KD and hypoxia for H3K36me2 (bottom), in control or sh*Kdm8*-expressing cells (F), and in 688M cells cultured in normoxia or hypoxia (G).

**Figure S6, related to Figure 6.**
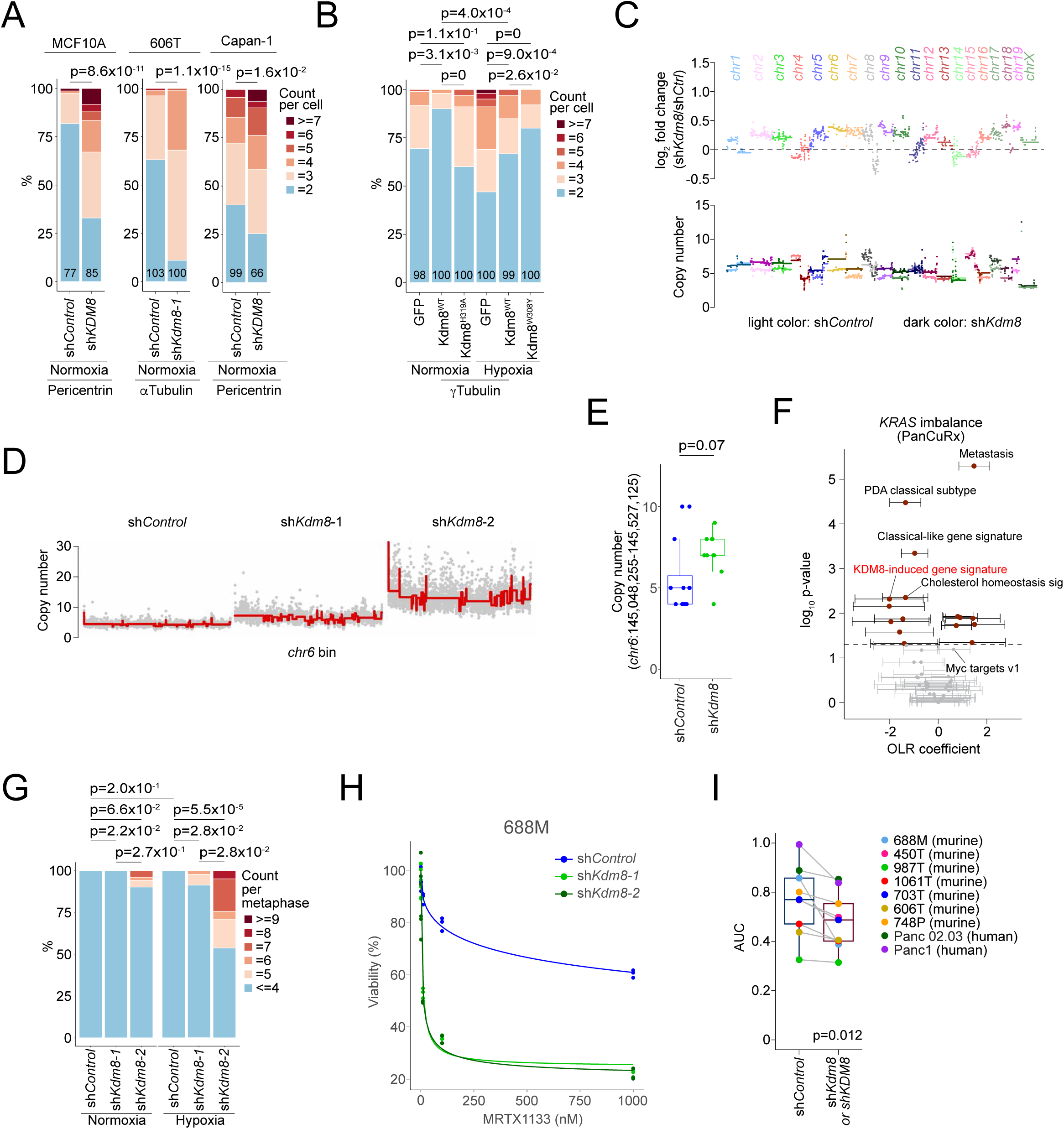
(A) Quantifications of anaphase polarities by Pericentrin or αTubulin immunofluorescence (IF) in indicated control and *Kdm8*/*KDM8* deficient PDA cells as in Figure 6E. Numbers in the plot indicate the number of cells quantified in each group. (B) Proportions of *Kdm8* knockout 688M cells re-expressing indicated Kdm8 variants cultured in normoxia or hypoxia with indicated anaphase polarities quantified by γTubulin IF. Numbers in the plot indicate the number of cells quantified in each group. (C) Single-cell whole genome sequencing (scWGS) showing log_2_ fold change (sh*Kdm8*/sh*Control*, top) and the copy number of genomes in control (n=12) and sh*Kdm8*-expressing 688M cells (n=11, bottom). (D) Bulk whole genome sequencing showing chromosome 6 copy-number alterations in control and 688M cells expressing two sh*Kdm8*s. (E) Copy numbers of the indicated genomic location on chromosome 6 (mm39) where *Kras* is located in control and sh*Kdm8*-expressing 688M cells using scWGS as in (C). p-value = 0.07 using two-sided Wilcoxon test. (F) Ordinal logistic regression (OLR) for the association between the *KRAS* allelic imbalance (wt, 0; balanced, 1; minor, 2; major, 3) and the GSEA hallmark transcriptomic programs, anatomic location (primary site, 0; liver metastasis, 1), PDA subtype (classical, 1; basal, 0), and KDM8-induced gene signature in the PanCuRx PDA cohort. Coefficients with the 2.5/97.5% confidence intervals (x-axis) and log_10_ converted p-values (y-axis) are shown. KDM8-induced gene signature is highlighted. Dotted line represents statistical significance (p=0.05). (G) Quantifications of FISH as in Figure 6I using a chromosome 18 centromeric probe (yellow, Figure 6H) in control and sh*Kdm8*-expressing 688M cells cultured in normoxia and hypoxia. Proportions of control and the sh*Kdm8* metaphase 688M cells cultured in normoxia or hypoxia are shown. (H) Percent viability of 688M cells expressing control or *Kdm8*-targeting shRNAs treated with indicated concentrations of Kras^G12D^ inhibitor MRTX1133. Results from three repeated experiments are shown. (I) Boxplot showing responses to MRTX1133 quantified by the area under the dose-response curve (AUC) for the indicated PDA cell lines. Significance was determined via paired student t-test following a Shapiro-Wilk test for normality.

**Table S1.** Survival and clinical features of disease progression in the PDA GEM model.

| Date of birth | Date of surgery | Date of death | Days | Surgical Procedure | Animal ID | Mouse genotype | Virus infused | Gender | Infused volume (uL) | PDA tumor | Splenomegaly | Ascites | Lung met # | Liver met # | Peritoneal met |
| --- | --- | --- | --- | --- | --- | --- | --- | --- | --- | --- | --- | --- | --- | --- | --- |
| 3/17/22 | 5/6/22 | 2/16/23 | 286 | Pancreatic ductal injection | 147 | PTC | AAV-U6-sgKdm8(J1)-KrasG12D(BC) | F | 80 | Y | Y | Y | 26 | 0 | Y |
| 3/17/22 | 5/6/22 | 11/19/22 | 197 | Pancreatic ductal injection | 305 | PTC | AAV-U6-sgKdm8(J1)-KrasG12D(BC) | F | 110 | N | NA | NA | NA | NA | NA |
| 3/17/22 | 5/11/22 | 2/27/23 | 292 | Pancreatic ductal injection | 306 | PTC | AAV-U6-sgKdm8(J1)-KrasG12D(BC) | M | 10 | N | NA | NA | NA | NA | NA |
| 3/17/22 | 5/11/22 | 2/27/23 | 292 | Pancreatic ductal injection | 142 | PTC | AAV-U6-sgKdm8(J1)-KrasG12D(BC) | M | 100 | N | NA | NA | NA | NA | NA |
| 3/17/22 | 5/11/22 | 2/28/23 | 293 | Pancreatic ductal injection | 307 | PTC | AAV-U6-sgKdm8(J1)-KrasG12D(BC) | F | 70 | Y | Y | Y | 47 | 0 | Y |
| 3/24/22 | 5/11/22 | 11/4/22 | 177 | Pancreatic ductal injection | 308 | PTC | AAV-U6-sgKdm8(J1)-KrasG12D(BC) | M | 80 | Y | Y | Y | 8 | NA | N |
| 5/23/22 | 6/29/22 | 2/8/23 | 224 | Pancreatic ductal injection | 214 | PTC | AAV-U6-sgKdm8(J1)-KrasG12D(BC) | M | 100 | Y | Y | Y | 109 | 12 | Y |
| 5/23/22 | 6/29/22 | 10/27/22 | 120 | Pancreatic ductal injection | 215 | PTC | AAV-U6-sgKdm8(J1)-KrasG12D(BC) | M | 90 | N | NA | NA | NA | NA | NA |
| 5/23/22 | 6/29/22 | 12/9/22 | 163 | Pancreatic ductal injection | 216 | PTC | AAV-U6-sgKdm8(J1)-KrasG12D(BC) | M | 90 | N | NA | NA | NA | NA | NA |
| 5/23/22 | 6/29/22 | 1/2/23 | 187 | Pancreatic ductal injection | 217 | PTC | AAV-U6-sgKdm8(J1)-KrasG12D(BC) | M | 100 | Y | N | Y | 51 | 18 | Y |
| 5/23/22 | 6/29/22 | 1/27/23 | 212 | Pancreatic ductal injection | 218 | PTC | AAV-U6-sgKdm8(J1)-KrasG12D(BC) | M | 120 | Y | Y | Y | 39 | 4 | Y |
| 5/23/22 | 6/29/22 | 10/21/22 | 114 | Pancreatic ductal injection | 219 | PTC | AAV-U6-sgKdm8(J1)-KrasG12D(BC) | M | 130 | N | NA | NA | NA | NA | NA |
| 8/29/22 | 10/6/22 | 1/24/23 | 110 | Pancreatic ductal injection | 252 | PTC | AAV-U6-sgKdm8(J1)-KrasG12D(BC) | F | 50 | Y | Y | Y | 60 | 2 | Y |
| 8/29/22 | 10/6/22 | 2/18/23 | 135 | Pancreatic ductal injection | 253 | PTC | AAV-U6-sgKdm8(J1)-KrasG12D(BC) | M | 50 | Y | Y | Y | 25 | 1 | Y |
| 8/29/22 | 10/6/22 | 2/3/23 | 120 | Pancreatic ductal injection | 254 | PTC | AAV-U6-sgKdm8(J1)-KrasG12D(BC) | M | 50 | Y | Y | N | 20 | 0 | Y |
| 1/12/23 | 2/22/23 | 7/17/23 | 145 | Pancreatic ductal injection | 378 | PTC | AAV-U6-sgKdm8(J1)-KrasG12D(BC) | M | 100 | N | NA | NA | NA | NA | NA |
| 1/12/23 | 2/22/23 | 7/17/23 | 145 | Pancreatic ductal injection | 379 | PTC | AAV-U6-sgKdm8(J1)-KrasG12D(BC) | M | 50 | N | NA | NA | NA | NA | NA |
| 1/12/23 | 2/22/23 | 6/21/23 | 119 | Pancreatic ductal injection | 501 | PTC | AAV-U6-sgKdm8(J1)-KrasG12D(BC) | M | 50 | Y | N | Y | 32 | 2 | Y |
| 3/28/23 | 5/22/23 | 4/4/24 | 318 | Pancreatic ductal injection | 449 | PTC | AAV-U6-sgNT-KrasG12D | M | 40 | Y | NA | N | 6 | 0 | N |
| 3/28/23 | 5/22/23 | 12/5/23 | 197 | Pancreatic ductal injection | 450 | PTC | AAV-U6-sgNT-KrasG12D | M | 280 | Y | NA | Y | 8 | 0 | N |
| 4/3/23 | 5/22/23 | 11/17/23 | 179 | Pancreatic ductal injection | 488 | PTC | AAV-U6-sgNT-KrasG12D | M | 80 | Y | Y | Y | 15 | 0 | Y |
| 4/3/23 | 5/22/23 | 2/15/24 | 269 | Pancreatic ductal injection | 491 | PTC | AAV-U6-sgNT-KrasG12D | M | 30 | Y | NA | N | 18 | 0 | N |
| 4/22/23 | 5/22/23 | 12/26/23 | 218 | Pancreatic ductal injection | 606 | PTC | AAV-U6-sgNT-KrasG12D | M | 70 | Y | Y | Y | 7 | 0 | N |
| 4/22/23 | 5/22/23 | 1/2/24 | 225 | Pancreatic ductal injection | 608 | PTC | AAV-U6-sgNT-KrasG12D | M | 60 | Y | Y | Y | 24 | 0 | N |
| 4/22/23 | 5/22/23 | 11/28/23 | 190 | Pancreatic ductal injection | 609 | PTC | AAV-U6-sgNT-KrasG12D | M | 120 | Y | Y | Y | 12 | 0 | N |
| 4/22/23 | 5/22/23 | 4/4/24 | 318 | Pancreatic ductal injection | 612 | PTC | AAV-U6-sgNT-KrasG12D | M | 25 | Y | Y | Y | 88 | 0 | N |
| 4/22/23 | 5/22/23 | 11/7/23 | 169 | Pancreatic ductal injection | 613 | PTC | AAV-U6-sgNT-KrasG12D | M | 40 | Y | NA | NA | NA | NA | NA |
| 10/24/24 | 12/10/24 | 4/23/25 | 134 | Pancreatic ductal injection | 2445 | PTC | AAV-U6-sgNT-KrasG12D | F | 20 | Y | Y | Y | 18 | 0 | Y |
| 10/24/24 | 12/10/24 | 4/25/25 | 136 | Pancreatic ductal injection | 2446 | PTC | AAV-U6-sgNT-KrasG12D | F | 50 | Y | Y | Y | 22 | 0 | N |
| 10/30/24 | 12/10/24 | 4/23/25 | 134 | Pancreatic ductal injection | 2429 | PTC | AAV-U6-sgNT-KrasG12D | F | 50 | Y | Y | Y | 1 | 0 | N |
| 10/31/24 | 12/10/24 | 4/7/25 | 118 | Pancreatic ductal injection | 2487 | PTC | AAV-U6-sgNT-KrasG12D | F | 70 | Y | Y | N | 77 | 0 | N |

